# Desmoglein-2 deficiency results in cardiac dysfunction by compromising both Z-disc- and intercalated disc-mediated mechanotransduction

**DOI:** 10.1101/2025.10.03.680335

**Authors:** Maicon Landim-Vieira, Vivek P. Jani, Emily Shiel, Hosna Rastegarpouyani, Morgan Engel, Flair Paradine, Ronnie Chastain, Waleed Farra, Weikang Ma, Christopher Toepfer, P. Bryant Chase, David A. Kass, J. Renato Pinto, Stephen P. Chelko

**Author notes:** Department of Biology, Illinois Institute of Technology, Chicago, IL, USA. **Co-corresponding authors:** Stephen Chelko, PhD, FACC, FHRS; Maicon Land im-Vieira, PhD mland.

## Abstract

Desmoglein-2 (DSG2), a critical component of the cardiac desmosome and located at the cardiomyocyte-cardiomyocyte intercalated disc, is essential for cell-cell adhesion, cardiomyocyte mechanical stability, and electrical coupling between cells. However, its relative contribution in maintaining cardiac function at the sarcomere level remains unclear. Using 4-week-old (adolescent) and 16-week-old (adult) homozygous knock-in *Dsg2*-mutant (*Dsg2*^mut/mut^) mice, we found that loss of DSG2 leads to early onset chamber- and age-dependent cardiac dysfunction driven by Z-disc structural defects and increased myosin detachment rate. Interestingly, Ca²⁺-activated force was markedly reduced in adolescent *Dsg2*^mut/mut^ permeabilized left ventricular cardiac muscle bundles but preserved in permeabilized isolated cardiomyocytes. This disparity demonstrates that DSG2 is not only crucial for mechanical coupling between cardiomyocytes but also for force transmission within and between sarcomeres, revealing a novel role for DSG2 in maintaining contractile integrity at both the cellular and tissue levels.

## INTRODUCTION

The cardiac desmosome is an abundant protein complex in the heart and found at the cardiomyocyte-cardiomyocyte intercalated disc (ICD), where they anchor adjacent cardiomyocytes and facilitate the coordinated contraction of cardiac muscle (1, 2). Desmoglein-2 (DSG2) is a key constituent of the cardiac desmosome and sustains intercellular mechanical and structural integrity of the ICD, helps facilitate electrical propagation between cells through gap junction proteins, and preserves central intracellular signaling pathways (1, 3). *DSG2* is the second most prevalent desmosomal-linked gene in arrhythmogenic cardiomyopathy (ACM), and its pathogenic variants cause a more severe form of ACM with an increased risk of heart failure (4). Functionally, ACM is characterized by cardiac dysfunction, ventricular wall motion abnormalities and aneurysms, and malignant ventricular arrhythmias; pathologically it is characterized by abnormal distribution of ICD proteins and myocardial inflammation and fibrosis (5, 6). Prior work demonstrated desmosomal dysfunction leads to Wnt/β-catenin suppression (7) and NFĸB activation (8), leading to immune cell chemotaxis in hearts of ACM subjects (9). These prior studies elucidated a central signaling pathway that contributes to myocardial inflammation in ACM. To date, little is known about the extent by which desmoglein-2 (DSG2) plays a role in cardiac contractile function at the sarcomere level, and the desmosome–sarcomere relation is not well defined. This gap in understanding may come from a historical focus on ICD alterations and arrhythmogenesis, at which point it may be difficult to isolate the effect of such alterations on sarcomere function. What is known, however, is that deletion of another desmosomal protein, desmoplakin (DSP), disrupts Z-disc structure (10).

In this study, we tested the hypothesis that deletion of DSG2 induces contractile dysfunction by disruption of the Z-disc, similarly observed with in DSP-deficient mice (10). To study the role of DSG2 in cardiac muscle regulation, we utilized a loss-of-function, homozygous knock-in *Dsg2*-mutant (*Dsg2*^mut/mut^) mouse model of ACM at 4-(adolescent) and 16-weeks of age (adult) *Dsg2*^mut/mut^ mice display ventricular dysfunction, increased ectopic burden, and myocardial fibrosis and inflammation by early adulthood (7–9, 11); replicating key disease characteristics observed in patients with ACM. Here, we show that the contractile effects of DSG2 deficiency are not uniformly manifested across the ventricles and stages of life; rather, contractile function appears to be both age- and chamber-specific during disease progression. Contractile dysfunction appears early, preceding overt morphological changes, and is likely driven by compromised Z-disc integrity, leading to reduced cross-bridge stiffness, and accelerated myosin detachment. Furthermore, given that DSG2 resides at the cell–cell junctions, we aimed to investigate its role in sarcomere structure and function using both permeabilized single cardiomyocytes and cardiac multicellular muscle bundles (CMBs). Our findings suggest that this dysfunction is not due to a loss of sarcomere function but rather a disruption in Z-disc- and ICD-mediated mechanotransduction.

## RESULTS

### Deficiency of DSG2 results in early presentation of cardiac dysfunction in adolescent *Dsg2*^mut/mut^ mice

Previous studies showed that adult *Dsg2*^mut/mut^ mice exhibit reduced cardiac function and overt cardiac remodeling (7, 11). To investigate whether deficiency of DSG2 results in early cardiac dysfunction, we evaluated echocardiographic (Echo) parameters in adolescent *Dsg2*^mut/mut^ mice and age-matched WT controls. *Dsg2*^mut/mut^ mice displayed heterogeneous wall motion abnormalities, including hypokinesia, dyskinesia, and areas of hyperkinesia in the LV anterior and posterior free walls (Fig.1a and Suppl. Table 1) as well as reduced LV fractional shortening, ejection fraction, stroke volume, and by extension, cardiac output (Fig.1c-d and Suppl. Table 1). Furthermore, *Dsg2*^mut/mut^ mice also showed reduced mitral valve early peak flow velocity/atrial peak flow velocity ratio (Fig.1b and Suppl. Table 1). No observable changes were noted in ventricular chamber size, wall thickness, or mass versus controls (Suppl. Table 1). Histopathological analysis of *Dsg2*^mut/mut^ mice detected no apparent signs of cardiac pathological remodeling and fibrosis (Fig. 1e-i), despite later presenting with pronounced myocardial fibrosis in adult *Dsg2*^mut/mut^ mice (11). Taken together, these data show that DSG2 deficiency results in the early cardiac dysfunction without apparent changes in heart morphology and fibrotic lesions.

**FIGURE 1.**
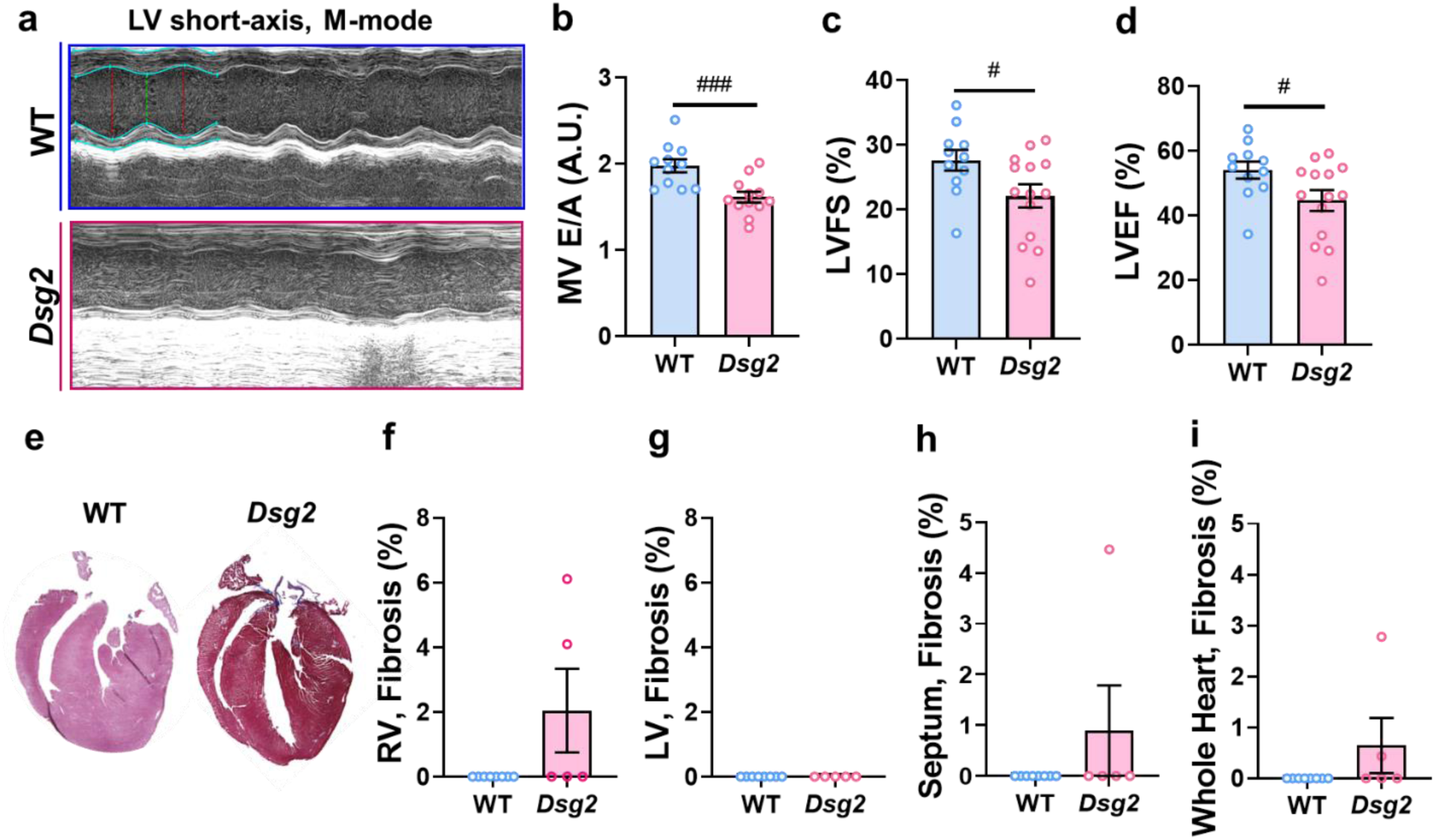
Figure legend 1. Absence of DSG2 results in early onset of cardiac dysfunction. **a)** Representative short-axis, m-mode echocardiograms from adolescent WT and *Dsg2*^mut/mut^ mice. **b)** Mitral valve early peak flow velocity – atrial peak flow velocity ratio (MV E/A). Percent left ventricular **c)** fraction shortening and **d)** ejection fraction (%LVFS and %LVEF, respectively). **e)** Representative cardiac tissue slices stained with Masson’s Trichrome for fibrotic lesions from adolescent WT mice and *Dsg2*^mut/mut^. Percent fibrosis of **f)** RV and **g)** LV free wall, **h)** septum, and **i)** whole heart. Data are shown as mean ± SE. Statistical significance was assessed by unpaired Student *t*-test. ^#^ P < 0.05, ^###^ P < 0.001 for adolescent *Dsg2*^mut/mut^ vs. WT mice. Number of independent animals per group were utilized: ECHO, WT n = 11 and *Dsg2*^mut/mut^ n = 7-9; and fibrosis, WT n = 8 and *Dsg2*^mut/mut^ n = 5.

### Loss of DSG2 impairs cardiac contractility in an age- and chamber-dependent manner

In adolescent *Dsg2*^mut/mut^ mice, LV but not RV CMBs exhibited a ∼63% reduction in maximum steady-state isometric force compared to WT mice (Fig. 2a and f, and Suppl. Table 2). In the RV, there was a marked rise in force at multiple Ca²⁺ levels (including systolic) in adult versus adolescent WT mice, while the LV showed a slight increase only at supraphysiological levels of Ca²⁺ (pCa 4.5). This behavior was lost in *Dsg2*^mut/mut^ mice (Fig. 2a and f, and Suppl. Table 2). Myofilament Ca²⁺-sensitivity, as assessed by the Ca²⁺ needed to achieve 50% of peak force, was increased in adolescent *Dsg2*^mut/mut^ LV CMBs but reduced in adult *Dsg2*^mut/mut^ CMBs. No such differences were observed in the RV (Fig. 2b and g, and Suppl. Table 2).

**FIGURE 2.**
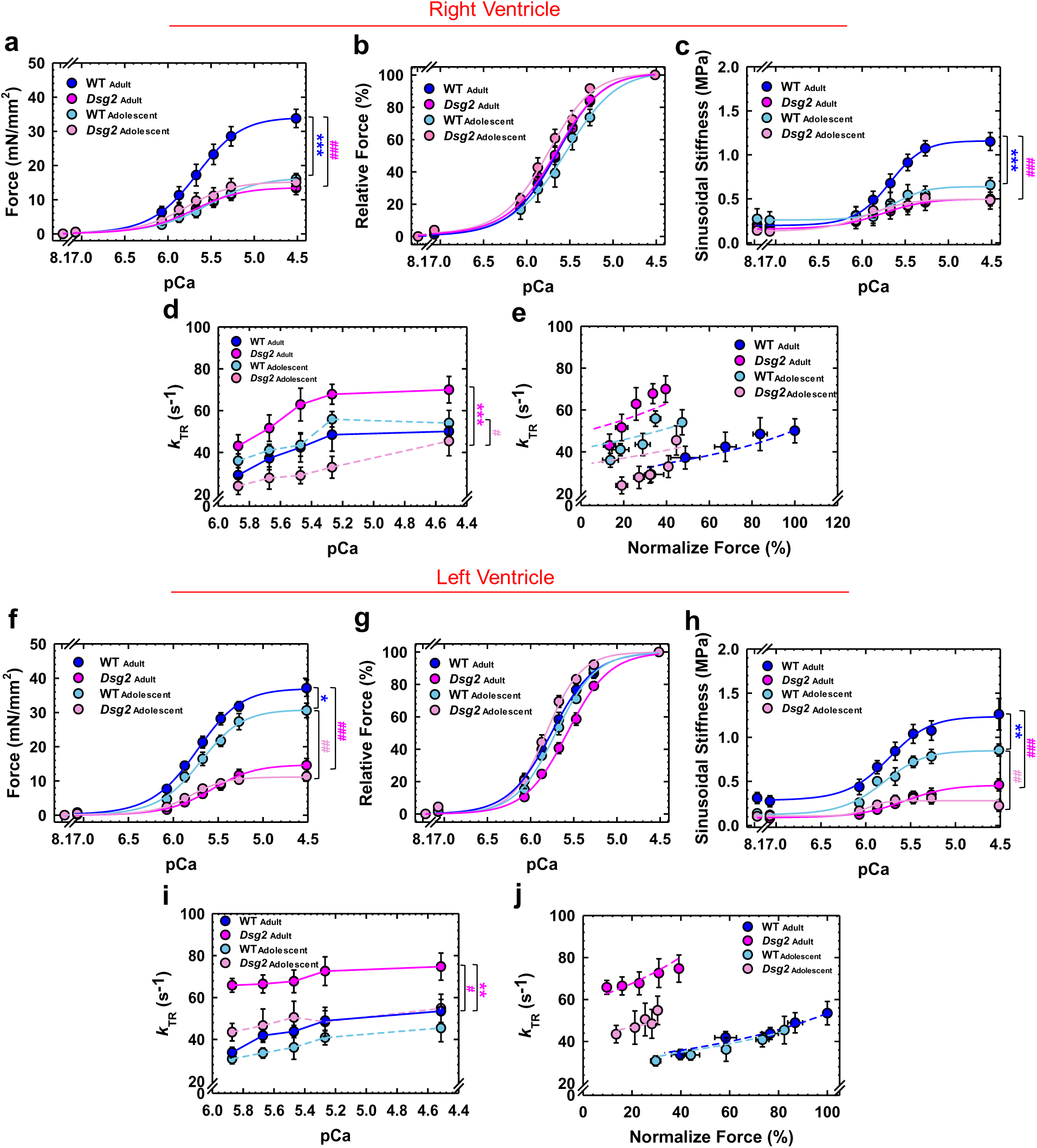
Figure legend 2. Altered sarcomere contractile parameters as a result of DSG2 deficiency. **a)** Steady-state isometric force levels normalized to the cross-sectional area of the cardiac muscle preparations. **b)** Relative steady-state isometric force as a function of Ca^2+^. The force values were normalized to the maximal steady-state isometric force in the same preparation. Ca^2+^-dependence of **c)** steady-state sinusoidal stiffness and **d)** kinetics of tension redevelopment (*k*_TR_). **e)** Normalized steady-state isometric force vs. *k*_TR_. All forces were normalized by the average force generated by adult WT. Dashed lines were obtained from fits to the 3-state model (Table 1). **f)** Steady-state isometric force levels normalized to the cross-sectional area of the cardiac muscle preparations. **g)** Relative steady-state isometric force as a function of Ca^2+^. The force values were normalized to the maximal steady-state isometric force in the same preparation. Ca^2+^-dependence of **h)** steady-state sinusoidal stiffness and **i)** kinetics of tension redevelopment (*k*_TR_). **j)** Normalized steady-state isometric force vs. *k*_TR_. All forces were normalized by the average force generated by adult WT. Dashed lines were obtained from fits to the 3-state model (Table 1). Data are shown as mean ± SE. Statistical significance was assessed by two-way ANOVA followed by Tukey’s or Bonferroni’s multiple comparison test: **a)** ^###^ P < 0.001 for adult *Dsg2*^mut/mut^ vs. WT at pCa 5.6 – pCa 4.5; *** P < 0.001 for adolescent WT vs. adult WT at pCa 5.6 – pCa 4.5. **c)** ^###^ P < 0.001 for adult *Dsg2*^mut/mut^ vs. WT at pCa 5.6 – pCa 4.5; *** P < 0.001 for adolescent WT vs. adult WT at pCa 5.4 – pCa 4.5. **d)** ^#^ P < 0.05 for neonatal *Dsg2*^mut/mut^ vs. WT at pCa 5.2; *** P < 0.001 for adolescent *Dsg2*^mut/mut^ vs. adult *Dsg2*^mut/mut^ at pCa 5.6 – pCa 4.5. **f)** * P < 0.05 for adolescent WT vs. adult WT at pCa 5.6, 5.4, and 4.5; ^###^ P < 0.001 for adult *Dsg2*^mut/mut^ vs. WT at pCa 6 – pCa 4.5; ## P < 0.01 for neonatal *Dsg2*^mut/mut^ vs. WT at pCa 5.8 – pCa 4.5. **h)** ^##^ P < 0.01 for adolescent *Dsg2*^mut/mut^ vs. WT at pCa 5.8 – pCa 4.5; ^###^ P < 0.001 for adult *Dsg2*^mut/mut^ vs. WT at pCa 6 – pCa 4.5; ** P < 0.01 for adolescent WT vs. adult WT at pCa 5.6 – pCa 4.5. **i) ^##^** P < 0.05 for adult *Dsg2*^mut/mut^ vs. WT at pCa 5.8 – pCa 5.2; * P < 0.05 for adolescent *Dsg2*^mut/mut^ vs. adult *Dsg2*^mut/mut^ at pCa 5.2. Number of independent animals per group: adolescent WT: 5-7 and adult WT:6-8; adolescent *Dsg2*^mut/mut^: 4-7 and adult *Dsg2*^mut/mut^: 6-9.

Next, we sought to identify the biophysical mechanism by which force is reduced. One important series of measurements involves assessment of crossbridge cycling kinetics, which, when impaired, result in reduced force development by reduction of the duty ratio. We assessed this via measuring the sinusoidal stiffness (SS) and kinetics of tension redevelopment (*k*_TR_) in CMBs. Our data revealed a decrease in SS that closely parallels the pattern of force decline observed in *Dsg2*^mut/mut^ mice. This reduction initially manifests in the LV of adolescent *Dsg2*^mut/mut^ mice and progressively becomes more pronounced in both ventricles over time (Fig. 2c and h, Ext. Fig.1a and c, and Suppl. Table 2). *k*_TR_ was significantly slower in adolescent *Dsg2*^mut/mut^ RVs compared to WT RVs, yet a faster *k*_TR_ was observed in adult RVs from *Dsg2*^mut/mut^ mice vs WT counterparts. In contrast, the LVs of *Dsg2*^mut/mut^ mice exhibited faster *k*_TR_ in both adolescent and adult stages as compared to their respective age-matched WT controls (Fig. 2d and i, Ext. Fig.1b and d, and Suppl. Table 2). Importantly, the absence of DSG2 had no effect on the Ca^2+^-dependence of cross-bridge cycling kinetics (Fig. 2d and i). A faster *k*_TR_ can be attributed to alterations in cross-bridge attachment (*f*) and detachment (*g*) rates. An increase in *k*_TR_ from an increase in *f* would be expected to elevate isometric force barring other compensatory mechanisms, while a *g-*dependent increase in k_TR_ would be expected to reduce generated tension. We further extended our analysis to mathematically estimate *f* and *g*, and the cardiac thin-filament *k*_OFF_ (Ca^2+^ OFF-rate constant) by employing a three-state model of cardiac muscle contraction (12). *k*_OFF_ was slightly reduced in both chambers from adolescent *Dsg2*^mut/mut^ mice, yet were not altered in adult *Dsg2*^mut/mut^ mice compared to WT controls (Fig. 2e and j, and Suppl. Table 3), which is in accordance with the belief that the thin filament has minimal contribution to declined contractility. In adolescent RVs, modeling predicted similar values for *f* and only modestly elevated *g* between groups, which could be attributed to the comparable force levels seen in the adolescent RVs (Fig.2 e, and Suppl. Table 3); whereas in adults, *Dsg2*^mut/mut^ RVs exhibited reduced *f* and increased *g*. Furthermore, modeling estimated that adolescent and adult *Dsg2*^mut/mut^ LVs exhibited a 2-fold reduction in *f* and a 2-fold increase in *g* (Fig. 2j, and Suppl. Table 3), which could explain the decline in steady-state isometric force levels.

Taken together, these data demonstrate that the absence of DSG2 exerts an early impact on force production by alterations in cross-bridge cycling kinetics solely within the LV, while the function of adolescent RVs remains preserved. However, over time, contraction in the RV becomes progressively dysfunctional.

### DSG2 deficiency affects cardiac sarcomere architecture, particularly Z-disc integrity

To investigate the root cause of altered cycling kinetics, we conducted a structural examination of the cardiac sarcomeres through electron microscopy (EM), which revealed notable differences in *Dsg2*^mut/mut^ mice compared to WT controls (Fig. 3a and b). Longitudinal sections of the RV from adult *Dsg2*^mut/mut^ mice revealed a decrease in sarcomere length (Fig. 3c) and an increase in I-band width (Fig. 3e) compared to WT mice. In contrast, longitudinal sections of the LV from adolescent *Dsg2*^mut/mut^ mice exhibited shorter sarcomeres (Fig. 3f) compared to WTs, a trend that persisted into adulthood. Interestingly, I-band width in the LV, which initially showed no changes, was later found to be reduced in adult *Dsg2*^mut/mut^ mice (Fig. 3h). No differences in A-band length were observed among cohorts (Fig. 3d and g).

**FIGURE 3.**
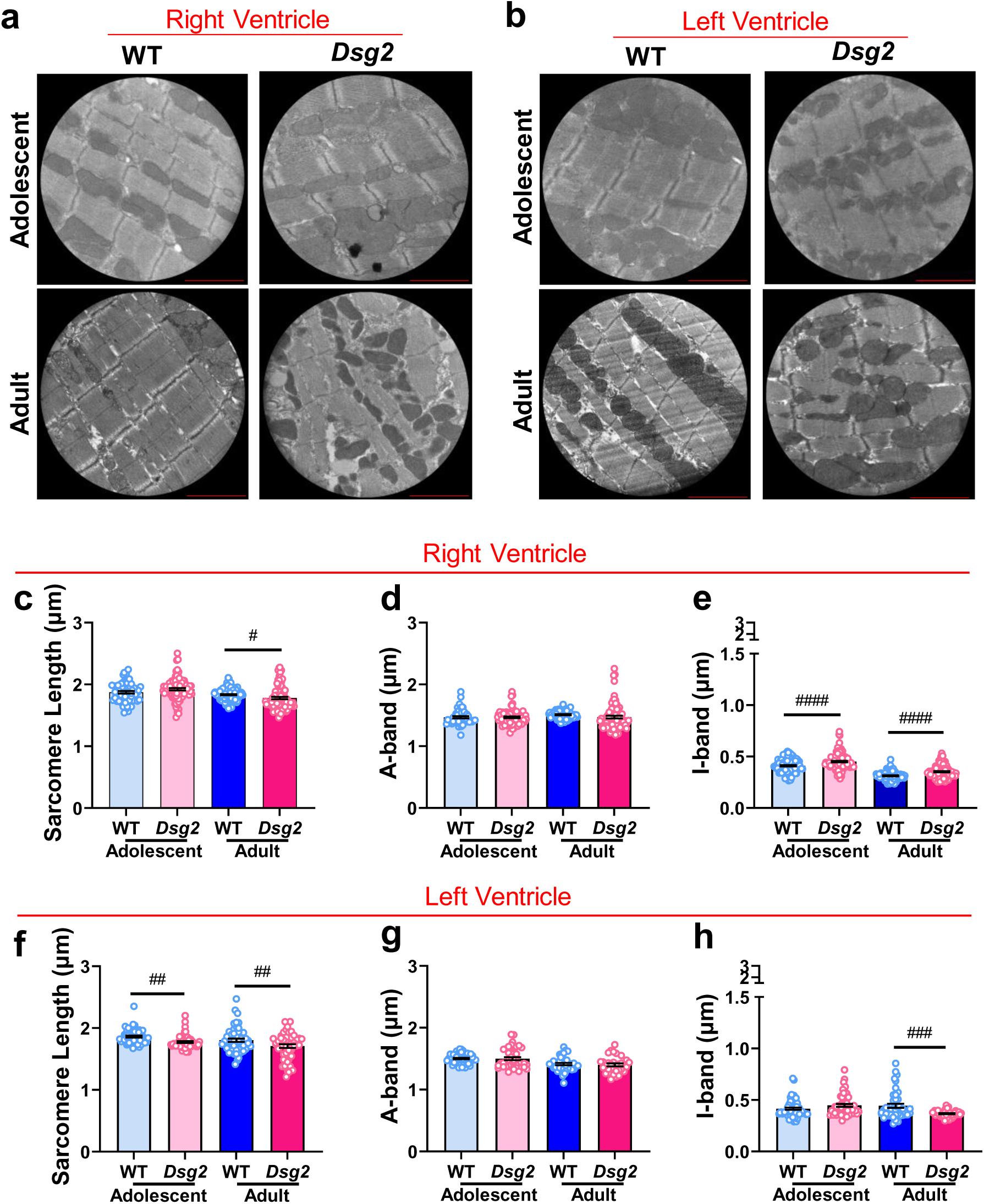
Figure legend 3. DSG2 deficiency affects the architecture of the cardiac sarcomere. **a-b** Representative images of the sarcomere structure in longitudinal sections of RV and LV free wall obtained from adolescent and adult WT and *Dsg2*^mut/mut^ mice. Measurements of both RV and LV sarcomere length (**c** and **f**), A-band (**d** and **g**), and I-band (**e** and **h**) are shown as mean ± SE. Statistical significance was assessed by two-way ANOVA followed by Bonferroni’s multiple comparisons test. ^#^ P < 0.05, ^##^ P < 0.01, and ^###^ P < 0.001 for *Dsg2*^mut/mut^ vs. WT mice. n = 3 independent animals per group were utilized. Scale bar = 2 µm. Sarcomere length: Adolescent RV, WT n = 63 and *Dsg2*^mut/mut^ n = 91; Adolescent LV, WT n = 57 and *Dsg2*^mut/mut^ n = 53; Adult RV, WT n = 108 and *Dsg2*^mut/mut^ n = 90; adult LV, WT n = 54 and *Dsg2*^mut/mut^ n = 51. A-band : Adolescent RV, WT n = 50 and *Dsg2*^mut/mut^ n = 73; Adolescent LV, WT n = 49 and *Dsg2*^mut/mut^ n = 43; Adult RV, WT n = 64 and *Dsg2*^mut/mut^ n = 85; adult LV, WT n = 39 and *Dsg2*^mut/mut^ n = 32. I-band : Adolescent RV, WT n = 101 and *Dsg2*^mut/mut^ n = 110; Adolescent LV, WT n = 61 and *Dsg2*^mut/mut^ n = 53; Adult RV, WT n = 89 and *Dsg2*^mut/mut^ n = 110; adult LV, WT n = 45 and *Dsg2*^mut/mut^ n = 45.

As such, we performed western blots (WB) to assess sarcomere and myofilament proteins and found that the levels of α-actinin-2 – a major component of the Z-disc – were decreased in both LV and RV from adult *Dsg2*^mut/mut^ mice (Fig.4a and e); but this was only observed in the LVs from adolescent *Dsg2*^mut/mut^ hearts. Further WB analyses of the myofilament proteins, cardiac troponin I (cTnI) and myosin-binding protein C (MyBP-C), along with their phosphorylation status, revealed a significant decrease in both protein content and phosphorylation levels primarily in the LVs of adult *Dsg2*^mut/mut^ mice (Suppl. Fig.1). Although LVs displayed reduced MyBP-C and cTnI content and phosphorylation levels, the ratio of phosphorylated to total protein were found to be unaffected (Suppl. Fig.1). Next, EM analysis revealed a significant reduction in Z-disc length (Fig.4b and f) and width (Fig.4c and g) in the LVs of both adolescent and adult *Dsg2*^mut/mut^ hearts. Intriguingly, the reduction in Z-disc length and width was observed exclusively in the LVs of adolescent *Dsg2*^mut/mut^ mice (Fig.4b and c, and f and g). Collectively, these findings strongly indicate that the absence of DSG2 exerts a pronounced influence on Z-disc integrity in an age- and chamber-specific manner, in which the LV is primarily affected at both adolescent and adult stages.

**FIGURE 4.**
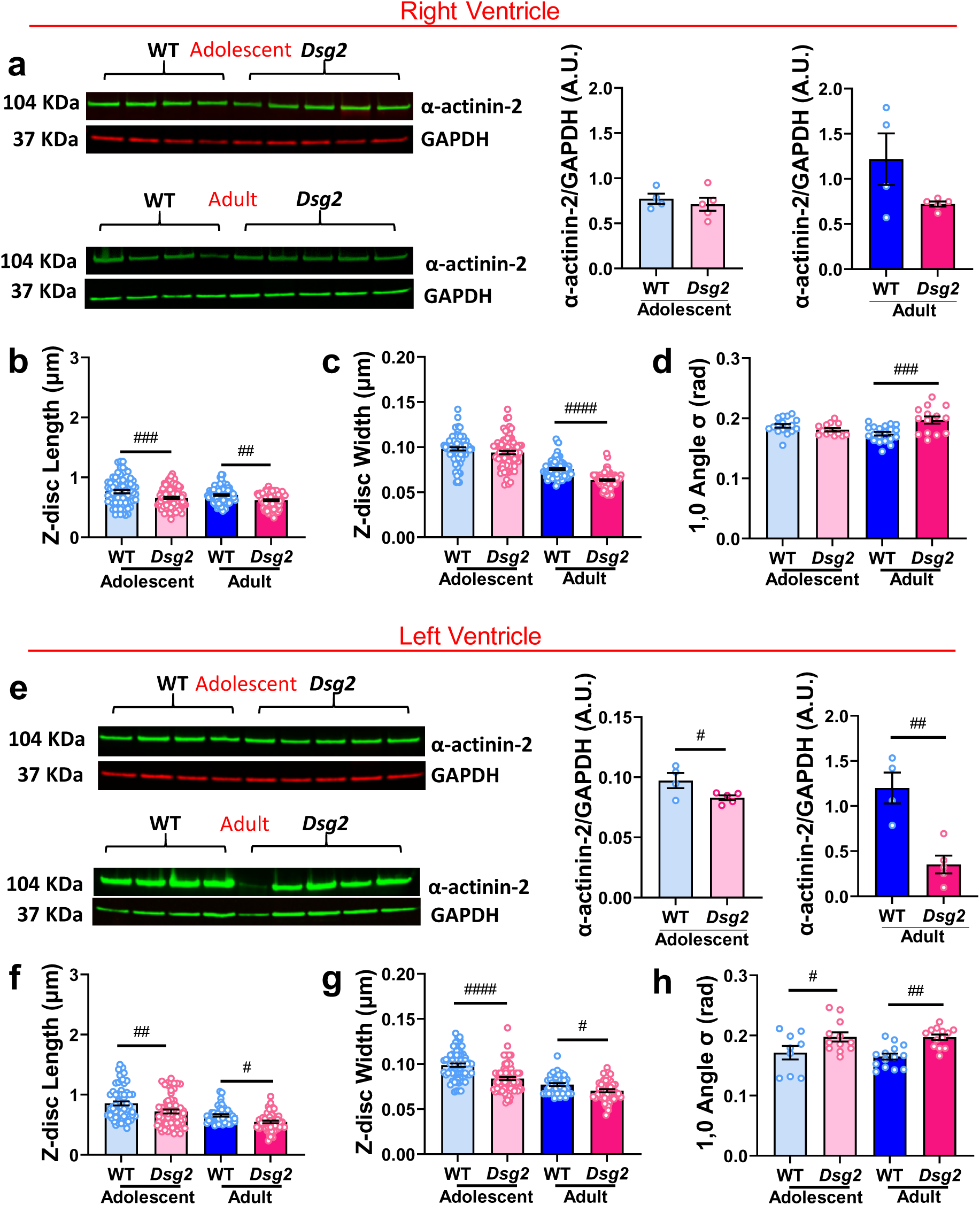
Figure legend 4. Lack of DSG2 compromises cardiac Z-disc integrity. **a)** Western blot analysis showing the protein levels of α-actinin-2 in the RV of adolescent and adult WT and *Dsg2*^mut/mut^. Quantification of α-actinin-2 relative to GAPDH in arbitrary units (A.U.). Measurements of Z-disc length **(b)** and width **(c)** in the RV of adolescent and adult WT and *Dsg2*^mut/mut^. **d)** The angular stand ard deviation of 1,0 equatorial reflections (angle σ) from longitudinal EM images of RV of adolescent and adult WT and *Dsg2*^mut/mut^ for myofilament disarray assessment. **e)** Western blot analysis showing the protein levels of α-actinin-2 in the LV of adolescent and adult WT and *Dsg2*^mut/mut^. Quantification of α-actinin-2 relative to GAPDH in arbitrary units (A.U.). Measurements of Z-disc length **(f)** and width **(g)** in the LV of adolescent and adult WT and *Dsg2*^mut/mut^. **h)** The angular stand ard deviation of 1,0 equatorial reflections (angle σ) from longitudinal EM images of LV of adolescent and adult WT and *Dsg2*^mut/mut^ for myofilament disarray assessment. Data are shown as mean ± SE. Western blot: statistical significance was assessed by unpaired Student *t*-test. ^#^ P < 0.05, ^##^ P < 0.001 between *Dsg2*^mut/mut^ vs. WT. *Dsg2*^mut/mut^, n = 5 and WT, n = 4 independent animals. Z-disc measurements: Statistical significance was assessed by two-way ANOVA followed by Bonferroni’s multiple comparisons test. # P < 0.05, ^##^ P < 0.01, and ^###^ P < 0.001 for *Dsg2*^mut/mut^ vs. WT mice. n = 3 independent animals per group were utilized. Z-disc length: Adolescent RV, WT n = 71 and *Dsg2*^mut/mut^ n = 101; Adolescent LV, WT n = 65 and *Dsg2*^mut/mut^ n = 63; Adult RV, WT n = 80 and *Dsg2*^mut/mut^ n = 78; adult LV, WT n = 48 and *Dsg2*^mut/mut^ n = 47. Z-disc width: Adolescent RV, WT n = 75 and *Dsg2*^mut/mut^ n = 83; Adolescent LV, WT n = 69 and *Dsg2*^mut/mut^ n = 75; Adult RV, WT n = 86 and *Dsg2*^mut/mut^ n = 73; adult LV, WT n = 42 and *Dsg2*^mut/mut^ n = 45. Myofilament disarray: Statistical significance was assessed by two-way ANOVA followed by Bonferroni’s multiple comparisons test. # P < 0.05, ^##^ P < 0.01, and ^###^ P < 0.001 for *Dsg2*^mut/mut^ vs. WT mice. n = 3 independent animals per group were utilized. Number of EM images: Adolescent RV, WT n = 17 and *Dsg2*^mut/mut^ n = 13; Adolescent LV, WT n = 9 and *Dsg2*^mut/mut^ n = 12; Adult RV, WT n = 20 and *Dsg2*^mut/mut^ n = 14; adult LV, WT n = 13 and *Dsg2*^mut/mut^ n = 13.

The Z-disc is a crucial structural component of the sarcomere, that maintains its integrity by anchoring the actin thin filaments, among other structures. In the absence of DSG2, the Z-discs were shorter and thinner in *Dsg2*^mut/mut^ mice (Fig.4), leading us to hypothesize that the myofilaments may not be properly aligned. Computational analysis of EM images revealed a misalignment of the myofilaments in *Dsg2*^mut/mut^ hearts (Fig.4d and h, and Ext. Fig. 2a). The latter originates in the LV of adolescent *Dsg2*^mut/mut^ mice and later becomes evident in both ventricles of *Dsg2*^mut/mut^ adults, paralleling the Z-disc changes mentioned above. Further computational analysis of the EM images revealed myofibrillar disarray, exclusively in *Dsg2*^mut/mut^ adult ventricles (Ext. Fig. 2b-d).

### Reduced myofilament lattice spacing in *Dsg2*^mut/mut^ CMBs

Given the observed alterations in Z-disc integrity and myofilament alignment (Figs.3 and 4), we further investigated whether *Dsg2*^mut/mut^ CMBs exhibit changes in the spatial organization of thick and thin filaments within the sarcomeres using small-angle X-ray diffraction (Fig.5a). Consistent with the observed myofilament disarray and altered Z-disc length and width in *Dsg2*^mut/mut^ mice, a reduction in lattice spacing was detected in the LV of adolescent *Dsg2*^mut/mut^ hearts (Fig. 5d), but not in the RV (Fig.5b). In adult *Dsg2*^mut/mut^ CMBs, the observed reduction in lattice spacing was present in both ventricles (Fig.5b and d), confirming that sarcomere structural alterations originate in the LV and later progress to the RV in murine *Dsg2*-linked ACM. Both RVs and LVs showed no changes in the equatorial intensity ratio (I_1,1_/I_1,0_) in relaxed CMBs, regardless of age (Fig.5c and e).

**FIGURE 5.**
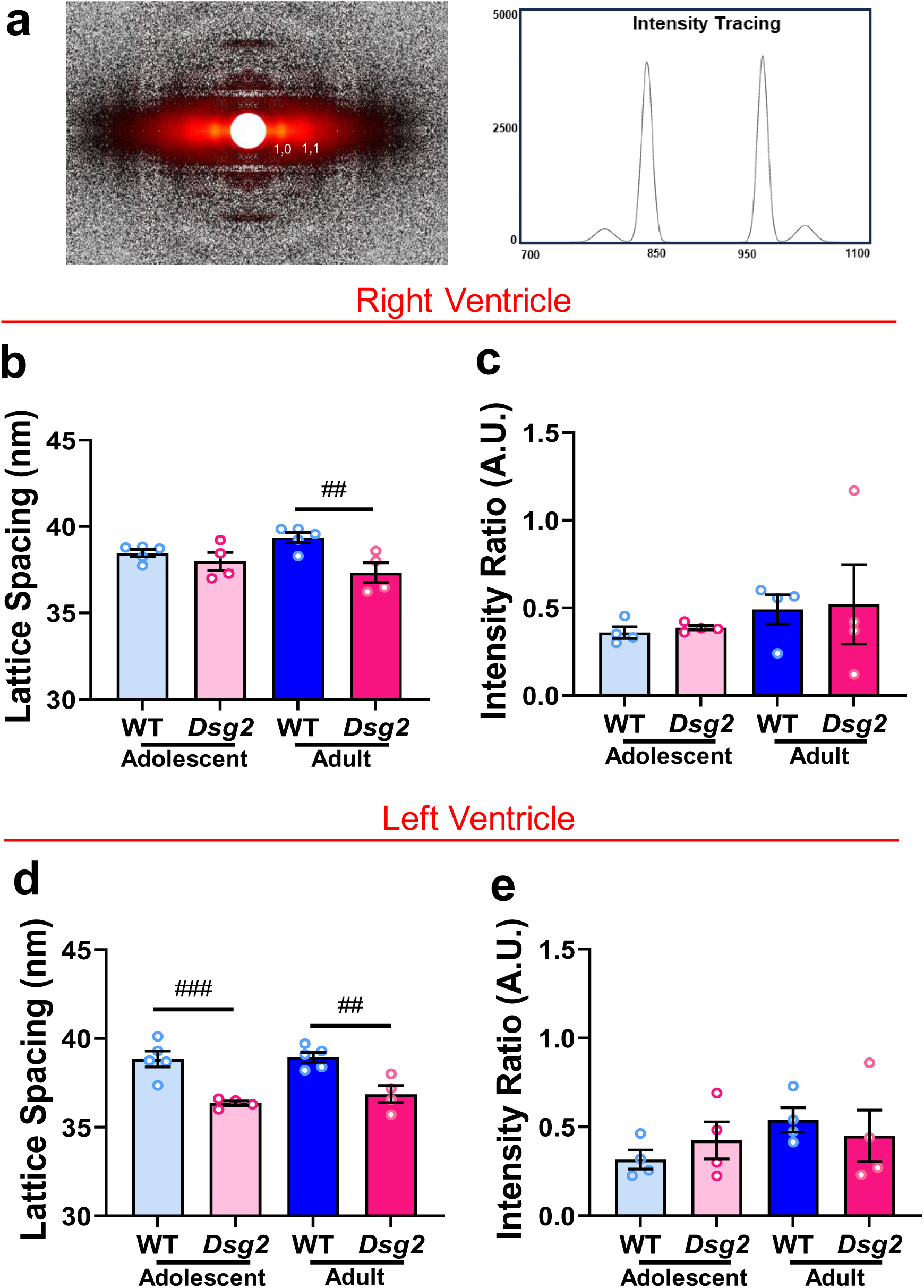
Figure legend 5. Small-angle X-ray diffraction patterns from *Dsg2*^mut/mut^ cardiac tissue revealed reduced myofilament lattice spacing. **a)** Representative small-angle X-ray diffraction pattern (left) and its equatorial reflection’s intensity tracing (right). Myofilament lattice spacing measurements obtained from RV **(b)** and LV **(d)** free wall of adolescent and adult WT and *Dsg2*^mut/mut^ mice. Intensity ratio of the equatorial reflections I_1,1_ and I_1,0_ obtained from RV **(c)** and LV **(e)** free wall of adolescent and adult WT and *Dsg2*^mut/mut^ mice. Data are shown as mean ± SE. Statistical significance was assessed by two-way ANOVA followed by Bonferroni’s multiple comparisons test. ^##^ P < 0.01 and ^###^ P < 0.001 *Dsg2*^mut/mut^ vs. WT mice. Number of independent animals used: WT, n = 4-5 and *Dsg2*^mut/mut^, n=4.

### Hearts from *Dsg2*^mut/mut^ mice exhibited a dysregulated SRX ↔ DRX equilibrium

To biochemically assess the impact due to the absence of DSG2 on the energetic states of myosin heads, we measured the percentage of myosin heads in the super-relaxed (SRX) and disordered-relaxed (DRX) states (13). During the adolescent stage, no alterations in the SRX ↔ DRX equilibrium was observed in *Dsg2*^mut/mut^ RVs (Fig.6a and b, and Ext. Fig. 3). However, in adult *Dsg2*^mut/mut^ mice, the percentage of RV myosin heads in the SRX state was significantly higher, and thus, the corresponding DRX state was far less (Fig.6a and b, and Ext. Fig. 3). Interestingly, this SRX ↔ DRX disequilibrium was already present in *Dsg2*^mut/mut^ LVs during the adolescent stage and persisted into adulthood (Fig.6c and d, and Ext. Fig. 3).

**FIGURE 6.**
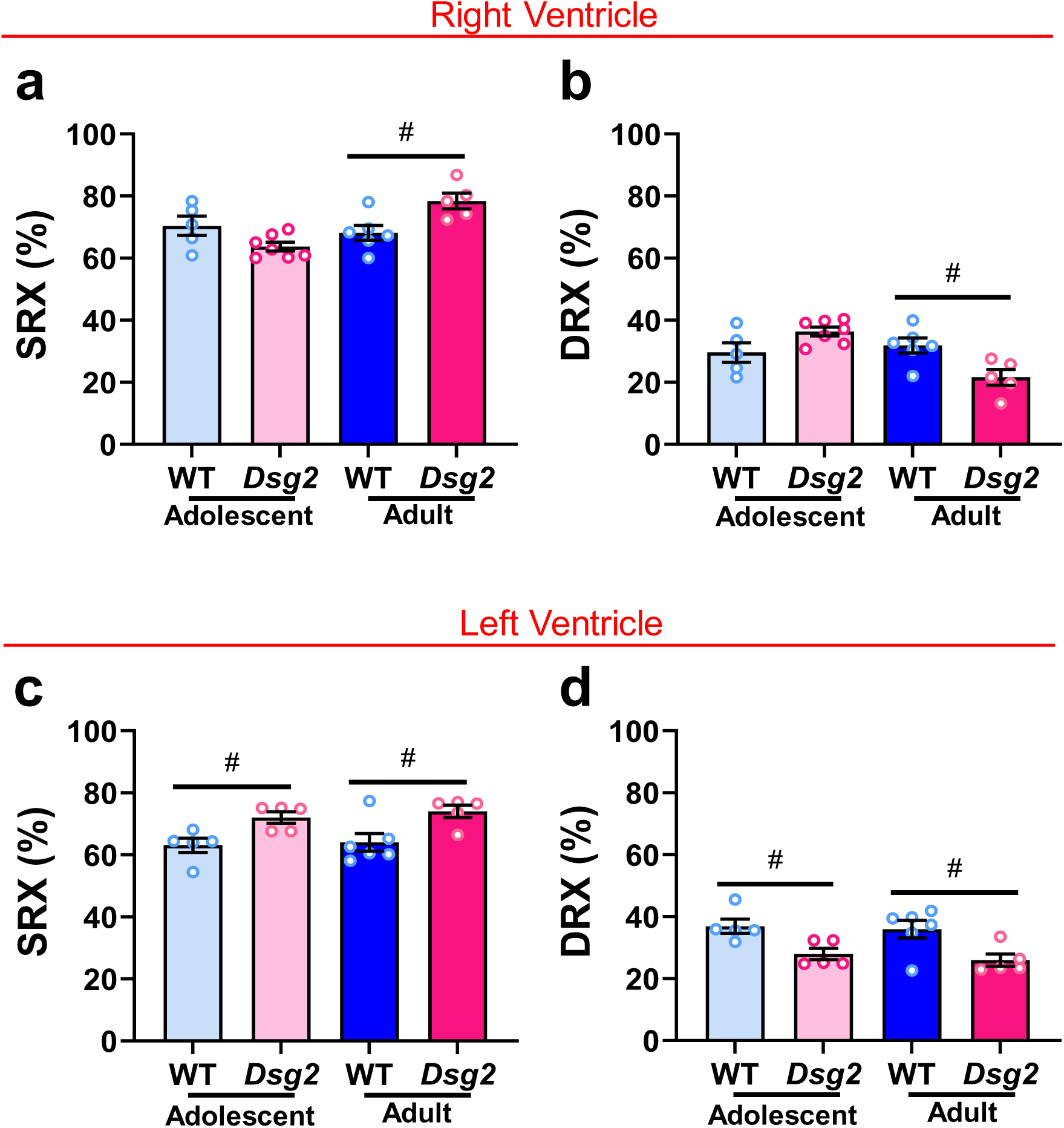
Figure legend 6. DSG2 deficiency increases the proportion of the SRX myosin population. Approximate percent of myosin heads in SRX (**a** and **c**) and DRX (**b** and **d**) was calculated using 0.6 as the correction factor to account for washout of non-myosin-bound Mant-ATP. Data are shown as mean ± SE. Statistical significance was assessed by two-way ANOVA followed by Bonferroni’s multiple comparisons test. ^#^ P < 0.05 *Dsg2*^mut/mut^ vs. WT mice. n = 5 independent animals per group were utilized. n = 5-6 and n = 5-7 technical replicates of WT and *Dsg2*^mut/mut^, respectively.

### Contractile dysfunction in adolescent *Dsg2*^mut/mut^ LVs was rescued by increased rate of crossbridge attachment

We show above, that *Dsg2*^mut/mut^ CMBs displayed alterations in sarcomere ultrastructure, myofilament alignment, lattice spacing, Z-disc integrity, and, most notably, contractile dysfunction, exclusively in the LV. We hypothesized that the absence of DSG2 compromises ICD stability, initiating a cascade of sarcomere alterations which can result in accelerated myosin detachment rate among other deficits. This instability can yield a number of deficits, one of which is a reduction in the mechanical force that can be sensed by this structure. If the reduction in contractile force is primarily from this mechanical instability, then anchoring the ends of a single cardiomyocyte to fixed posts, which can handle a theoretical infinite amount of force, should – in essence – overcome this deficit, by a compensatory acceleration in attachment rate. To test this, we first isolated single cardiomyocytes, which were then permeabilized and attached to a force transducer on one side and to a motor on the other, which served to anchor each end of a single cardiomyocyte such that the force generated was no longer dependent upon the structural stability of the ICD. Measurements of single-cell muscle mechanics showed no changes in steady-state isometric force levels in the adolescent RVs of *Dsg2*^mut/mut^ mice compared to WT controls, consistent with data from CMBs (Fig.7a). Additionally, no differences were observed in myofilament Ca^2+^-sensitivity (Fig.7b), thin filament cooperativity (Fig.7b), or *k*_TR_ (Fig.7c). As expected, there were no changes in either myosin attachment (Fig.7d) or detachment rates (Fig.7e). These findings confirm that DSG2 deficiency does not affect RV contractile function in this setting, corroborating our CMB results. In adolescent LV single cardiomyocytes, we observed no differences in steady-state isometric force levels (Fig.7f), myofilament Ca^2+^-sensitivity (Fig.7g), and thin filament cooperativity (Fig.7g) between *Dsg2*^mut/mut^ and WT mice. However, an increase in LV isolated myocyte *k*_TR_ was observed (Fig.7h), followed by increases in both myosin detachment, (Fig.7j) and, importantly, attachment rates (Fig.7i).

**FIGURE 7.**
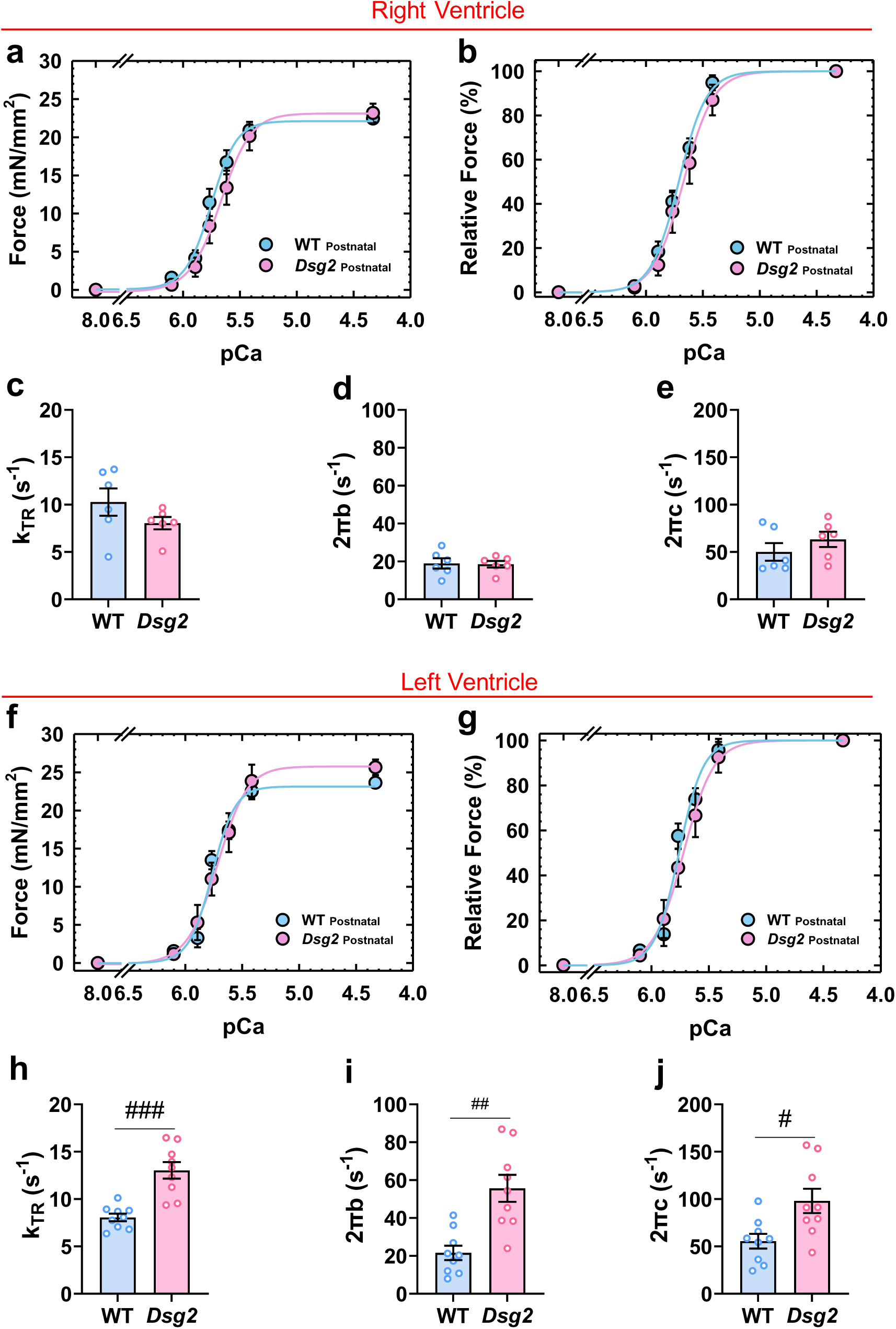
Figure legend 7. Cardiomyocytes from DSG2 deficient mice show preserved Ca^2+^-activated force and accelerated myosin detachment. **a** and **f)** Steady-state isometric force levels normalized to the cross-sectional area of the cardiac muscle preparations. **b** and **g)** Relative steady-state isometric force as a function of Ca^2+^. The force values were normalized to the maximal steady-state isometric force in the same preparation. **c** and **h)** maximal kinetics of tension redevelopment (*k*_TR_). **d** and **i**) 2πb correspond to attachment rate. **e** and **j**) 2πb correspond to detachment rate. Data are shown as mean ± SE. Statistical significance was assessed by two-way RM ANOVA (**a, b, f,** and **g**) and unpaired t test with Welch’s correction (**c-e** and **h-j**). ^#^ P < 0.05, ^##^ P < 0.01, and ^###^ P < 0.001 adolescent *Dsg2*^mut/mut^ vs. WT mice. n = 4 independent animals per group were utilized, and n = 7 and = 9-10 technical replicates of WT and *Dsg2*^mut/mut^, respectively.

## DISCUSSION

In this study, we utilized a robust mouse model of ACM (i.e., *Dsg2*^mut/mut^) to investigate the role of DSG2 on cardiac function at the sarcomere level. Both male and female *Dsg2*^mut/mut^ mice harbored a germline *Dsg2* loss-of-function gene variant resulting in a frameshift mutation, presence of four stop codons, and premature truncation due to nonsense-mediated mRNA decay (7). Our findings revealed a DSG2-dependent maturation process in the ventricles of WT mice, highlighting the critical role of DSG2 in maintaining long-term cardiac function. Conversely, in *Dsg2*^mut/mut^ mice, while the RV remained unscathed during early adolescence even in the absence of DSG2, it progressively failed to adapt to the ever-increasing mechanical stress and hemodynamic demands associated with aging, unlike the already compromised LVs observed in *Dsg2*^mut/mut^ mice. Notably, early contractile dysfunction precedes overt morphological changes, and is most likely the result of compromised Z-disc integrity, reduced cross-bridge stiffness, and accelerated myosin detachment. Our results suggest that contractile dysfunction is not simple due to a loss of sarcomere function, but rather the disruption of Z-disc- and ICD-mediated mechanotransduction. The latter, and unexpected finding, was confirmed in our isolated single cardiomyocyte preps.

Lamentably, ACM is frequently misdiagnosed as myocarditis (14) and the fact that ACM is plagued by incomplete penetrance and variable expressivity (15) complicate disease management; thus, confounding the elucidation of aberrant signaling mechanisms that contribute to disease onset and progression. It is therefore necessary to study the natural disease progression of ACM to better understand the underlying mechanisms of disease pathogenesis. A previous report diligently outlined the progression to heart failure in classical ACM, noting that arrhythmias typically emerge in the early stages of the disease, while progressive RV failure and LV dysfunction develops during its natural disease course (16). Here, our findings demonstrate that contractile dysfunction precedes overt myocardial fibrosis in *Dsg2*^mut/mut^ mice (Fig. 1). It has been reported in a different *Dsg2* mouse line, that myocyte necrosis, and myocardial calcification and fibrosis were present as early as 3-5 weeks of age (17). However, it should be noted that this *Dsg*2-linked mouse model of ACM harbored a cardiac-restricted *Dsg2* variant, unlike our model which harbors a germline *Dsg2* variant (i.e., mimicking human ACM).

By evaluating contractile parameters in *Dsg2*^mut/mut^ mice at 4 weeks of age, we show that contractile impairment occurs much earlier than previously reported in this *Dsg2* mutant mouse line (previously reported at 8 weeks of age). A previous study using transgenic mice with cardiac-specific overexpression of murine DSG2-N271S (human equivalent: DSG2-N266S) showed that SCD occurred by ∼3.5 weeks in 30% of mice that possessed the highest expression levels of DSG2-N271S (17). This is remarkably similar to the age when adolescent *Dsg2*^mut/mut^ mice exhibit early cardiac dysfunction and LV sarcomere-associated abnormalities (Figs. 1-6 and Suppl. Table 1). Gross morphological examination of hearts from DSG2-N271S mice younger than 2 weeks did not reveal morphological changes. However, by ∼4 weeks of age, DSG2-N271S mice displayed biventricular necrosis, increased heart weight, and whitish-fibrotic streaks in the subepicardial myocardium (17). Meanwhile, our adolescent *Dsg2*^mut/mut^ mice showed neither morphological nor heart weight changes compared to WT controls (Fig. 1 and Suppl. Table 1).

Maturation of the cardiac ventricles is a critical process that extends beyond embryonic development, continuing through the adolescent stage and into adulthood (18). A prior mouse study showed that approximately three weeks after birth, diastolic function matures, coinciding with the maturation of ventricular recoil and relaxation mechanisms (19). Here, we show that both ventricles in adult WT mice increased force levels compared to ventricles in adolescent WT mice; however, this finding was not observed in *Dsg2*^mut/mut^ mice (Fig. 2a and f, and Suppl. Table 2). This suggests that this process may be DSG2-dependent (or, at least, desmosome-dependent). During the adolescent stage, additional cardiac adaptations occur to continuously adjust to the persistent physiological demands, such as rapid cardiac growth (18, 20), increased heart rate (19), and rearrangement of the composition of the ICDs (21). It has been shown that *DSG2* variants can lead to disorganized ICDs, with indistinct or absent desmosomal-like plaques, frequent widening of the intercellular cleft, and partial or complete dissociation of cardiomyocytes at the ICD (22). Overall, the role of DSG2 deficiency in the impaired maturation of ventricles remains unclear.

Alpha-actinin-2 (encoded by the *ACTN2* gene) is a major component of the Z-disc that crosslinks actin filaments from adjacent sarcomeres, stabilizing sarcomere structure and ensuring efficient force transmission across myofibrils (23, 24). We showed *Dsg2*^mut/mut^ myocardium displays reduced alpha-actinin-2 levels, and thinner and shorter Z-discs (Fig. 4), which could affect cardiac contractility. Our findings contrast with previous reports in cardiomyocyte-specific desmoplakin (*Dsp*) knockout mice, which demonstrated widening of Z-discs (10). These differences emphasize the complexity of desmosomal-associated sarcomere regulation and suggest that individual desmosomal proteins may exert distinct effects on sarcomere stability. In addition, we show disarray of the of myofilaments in *Dsg2*^mut/mut^ myocardium. This aligns with findings reported in zebrafish with a cardiac-specific *Actn2* loss-of-function variant, where alpha-actinin-2 maintains precise lateral thin filament alignment within the Z-disc rather than abnormalities in its anchoring (25). Biopsies from a patient carrying an *ACTN2* variant showed structural and functional changes in the sarcomere, leading to a decline in cardiac contractile function (24). These findings suggest that weakened Z-discs fail to effectively transmit force across the sarcomere network, which resulted in a loss of contractile force, in part, due to individual sarcomeres that became mechanically decoupled from one another. We show here that the net effect of the latter is associated with desmosomal disruption and reduced contractility. Despite attributing the Z-disc abnormalities to the absence of DSG2, the precise mechanism by which the lack of DSG2 disrupts Z-disc remains unclear.

We propose two primary mechanisms responsible for cardiac dysfunction in *Dsg2*^mut/mut^ mice: (i) impaired Z-disc-mediated mechanotransduction and (ii) impaired ICD-mediated mechanotransduction. The Z-disc anchors thin filaments and provides minimal elastic resistance under relaxed conditions (26). During stretch activation, however, it becomes stiffer (i.e., increased elastance), distributing changes in thin filament strain that naturally occurs from cross-bridge cycling by redistributing these forces to neighboring thin filaments in the network (27). Here, we identified striking abnormalities in Z-disc structure accompanied by a reduction in alpha-actinin-2 (Fig. 4). We therefore propose that the thinning and shortening of the Z-disc prevents the Z-disc from increasing its elastance during contraction, resulting in an accelerated myosin detachment rate. Indeed, the latter has been similarly reported in an *ACTN2* variant (p.A868T) linked to hypertrophic cardiomyopathy (24). These sarcomeric abnormalities explain the concordant elevated myosin detachment rates observed in single cardiomyocytes and CMBs (Suppl. Table 3). Moreover, if this mechanism is true, then one would expect a reduction in sinusoidal stiffness, not from a reduction in the number of cross-bridges, but a reduction in the stiffness of individual cross-bridges. Indeed, sinusoidal stiffness was reduced in *Dsg2*^mut/mut^ CMBs (Fig. 2), while X-ray equatorial intensity ratio was preserved, suggesting equal amount of myosin heads moving towards the vicinity of the thin filament (Fig. 5).

One of our primary findings was the striking disparity between permeabilized single cardiomyocyte and CMB force generation in *Dsg2*^mut/mut^ experimental preps, the former being preserved and the latter reduced (Figs. 2 and 7). This discrepancy can be explained by impaired ICD-mediated mechanotransduction and Newtonian statics (Fig. 8). CMBs contain serial cardiomyocytes along with their ICDs. The maximum force generated by any single cardiomyocyte can be no more than what any one ICD can handle without rupture. This limitation is transduced through all neighboring myocytes via ICDs. In *Dsg2*^mut/mut^ mice, the deficiency of DSG2 likely results in weakening of the ICDs. It therefore follows that the force any cardiomyocyte produces within these CMBs will be less given the loss of ICD integrity. If this is a primary mechanism, then a system in which each end of a single cardiomyocyte is attached to two posts with theoretical infinite resistance should normalize the force generated. However, given the primary deficit in the Z-disc observed in our electron microscopy findings, this will occur in the context of aberrations in crossbridge cycling kinetics. Given that the structural integrity of attachments on either end of a single cardiomyocyte are now intact, the cardiomyocyte should produce normal force, which was found in this case (Fig. 7). The acceleration of myosin detachment is expected in this experimental prep given the effects that DSG2 absence had on Z-disc structure and function. Assuming the number of myosin heads are equivalent, and the stiffness of any single cross-bridge is identical to CMBs and individual cardiomyocytes, the only way this would be possible is by accelerating attachment, which was additionally observed (Figs. 2 and 7, and Table 1). We propose this to be a form of compensation by an isolated cardiomyocyte, and thus the cardiomyocyte achieves this by reducing myofilament lattice spacing (Fig. 5).

**FIGURE 8.**
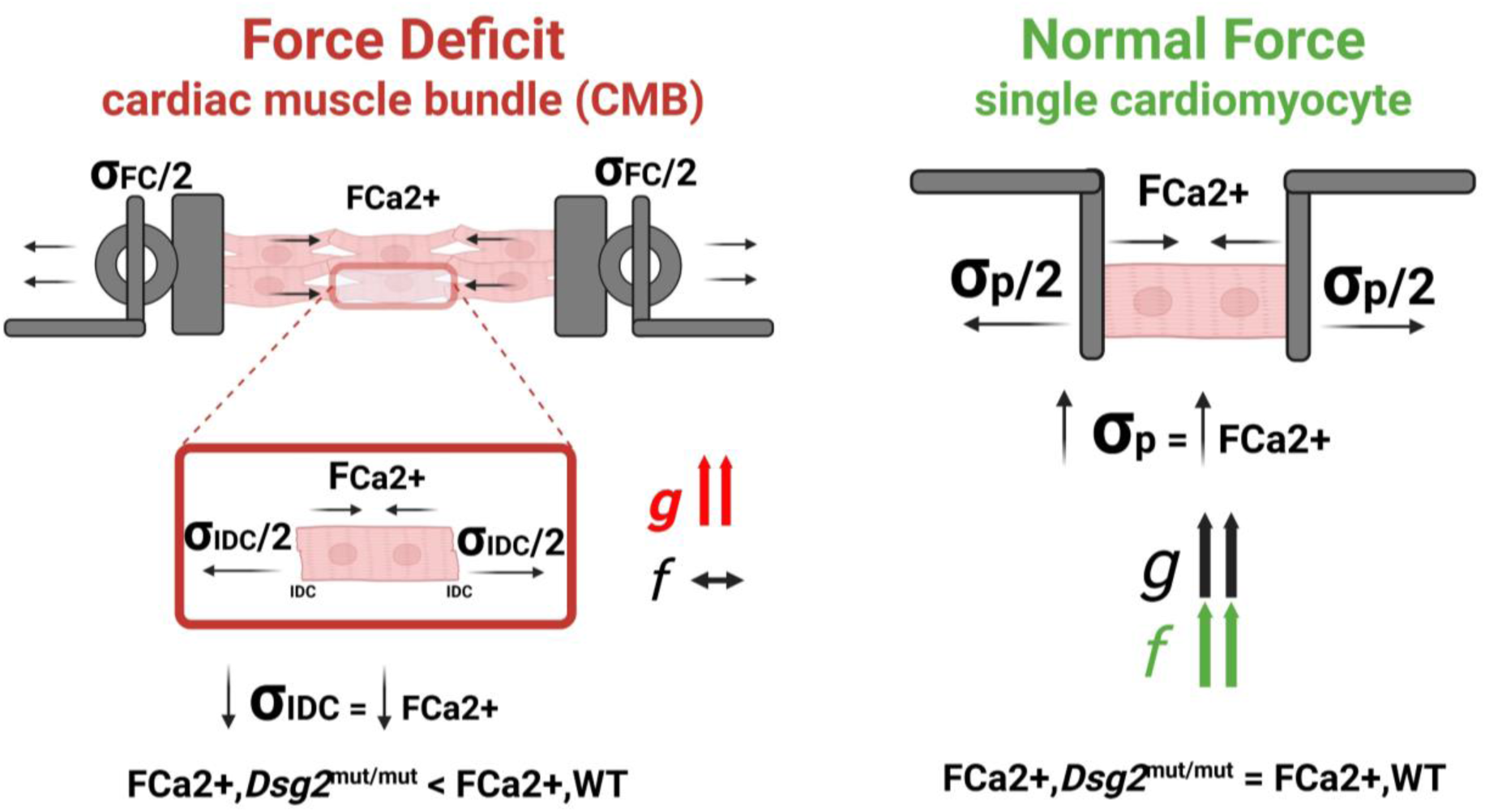
Figure legend 8. Proposed mechanism for force generation deficit in cardiac muscle bundles despite normal contractility in isolated cardiomyocytes from *Dsg2*^mut/mut^ hearts. Schematic representation comparing Ca²⁺-activated force production in cardiac muscle bundles (CMB, left) versus isolated single cardiomyocytes (right). In a multicellular context, the force generated by each cardiomyocyte is constrained by the mechanical integrity of intercalated discs (IDCs), which transduces force across neighboring cells. In *Dsg2*^mut/mut^ hearts, loss of DSG2 likely compromises IDC integrity, reducing IDC stress (σIDC) and elevating the detachment rate (*g*) while preserving the attachment rate (*f*). As a result, even if individual cardiomyocytes are intrinsically capable of generating force, overall bundle-level force (FCa²⁺) is diminished due to impaired force transmission through weakened IDCs. In contrast, isolated cardiomyocytes from *Dsg2*^mut/mut^ hearts generate normal force upon Ca²⁺ activation, equivalent to WT cardiomyocytes, despite an increased *g*. This suggests that when cardiomyocytes edges are stable (e.g., attached to experimental pins), effectively bypassing the defective IDCs, force output is restored. The elevated *g* must be compensated by an increase in *f*, assuming equivalent cross-bridge number and stiffness, to maintain normal force in single-cell preparations.

In conclusion, understand ing the age- and chamber-specific regulation of cardiac function by DSG2, along with the mechanotransduction defects associated with DSG2 alterations, is crucial for elucidating the pathogenesis of DSG2-linked ACM and related cardiomyopathies. This knowledge may also identify new therapeutic targets – such as strategies aimed at stabilizing sarcomere integrity or improving force transmission – to refine genotype–phenotype correlations in inherited cardiomyopathies, and enhance our understanding of mechano-signaling pathways that coordinate cellular adhesion and contraction. Ultimately, it could lead to a reclassification of DSG2-related disease as involving both junctional and contractile components, with important implications for diagnosis and treatment.

## METHODS

### Experimental Animals

All experiments conformed to the National Institutes of Health Guide for the Care and Use of Laboratory Animals (NIH publication no. 85–23, revised 1996), were approved by the Florida State University Animal Care and Use Committee, and conducted in accordance with appropriate ethical guidelines. Wild-type (WT) controls and homozygous Desmoglein-2 mutant (*Dsg2*^mut/mut^, or simply *Dsg2*) mice are of C57BL/6J background. *Dsg2*^mut/mut^ mice were generated as previously described (7). All experiments were performed utilizing cardiac tissue from both adolescent (4 weeks old; 4W) and adult (16 weeks old; 16W) age-matched female and male mice. 4W old mice were classified as adolescent, based on the prior study by Wang, Shuo et al. that correlated human with mouse ages, where 4W old mice are equivalent to a 1-year-old human (28). All mice were housed in a 12-hour dark/light cycle facility, in a climate-controlled room (18–22°C), and provided food and water *ad libitum*.

### Echocardiography

Echocardiography was performed using a Vevo F2 Visualsonic system (FUJIFILM VisualSonics, Toronto, ON, Canada) with an ultrahigh frequency (71 MHz) transducer. Echocardiographic images and videos were taken in accordance with the American Society of Echocardiography guidelines for animals (29). Both short- and long-axis images were acquired at the level of the papillary muscles with an ultrafast imaging sweep speed of >50,000 frames per second. All functional measurements were obtained using ≥3 echocardiographic images from each mouse then averaged to assess left ventricular function via percent ejection fraction (%EF), as previously described (7–9, 30).

### Histopathology

Excised hearts were washed in ice-cold 1X PBS (Phosphate-Buffered Saline), cut in half longitudinally, then incubated in 10% buffered formalin overnight. Following 24 hours in 10% buffered formalin, hearts were processed using a Tissue-Tek VIP6 AI vacuum infiltration processor then embedded in paraffin wax using a Tissue-Tek TEC6 embedder (Sakura Finetek USA, Inc., Torrance, CA). Formalin-fixed, paraffin-embedded blocks were cut at 5 μm, slices were mounted on positively charged slides, then underwent Masson’s Trichrome staining (Cat. No. HT15-1KT, Sigma Aldrich) using the Histo-Tek SL Slide Stainer (Sakura Finetek USA, Inc., Torrance, CA). Slides were imaged in brightfield using a Keyence BZ-X710 (Itasca, IL) microscope and percent myocardial fibrosis (%fibrosis) was determined as previously described (7–9, 30), using ImageJ version 1.8 software.

### Electron microscopy

Excised hearts were quickly and carefully soaked in PBS to make sure all the blood was washed away before starting the fixation process. To ensure optimal preservation of cellular structures, a multi-step fixation process was employed. Initially, the hearts were perfused with 0.1 M cacodylate buffer, followed by Karnovsky’s fixative containing 3% paraformaldehyde and 2.5% glutaraldehyde in 0.2 M sodium cacodylate buffer (pH 7.4). The heart samples were longitudinally sliced into halves, each containing both ventricles. The LV and RV free walls were then manually sectioned into 1×1×3 mm longitudinal blocks and left in fixative at 4°C overnight. Secondary fixation was performed using 1% OsO_4_ with 1.5% KFe for 30 minutes, followed by an additional fixation with 1% OsO_4_ for 60 minutes. Subsequently, the samples underwent overnight staining with 1% uranyl acetate and were subjected to a series of ethanol and propylene oxide dehydrations. The tissues were finally embedded in an epoxy resin embedding medium and incubated at 60°C for 48 hrs. Ultrathin sections (60 nm) were then obtained and placed on 200-mesh copper grids. Electron micrographs were captured using a Hitachi electron microscope (HT7800) operating at 100 kV. The selection of high-quality images with distinct sarcomeric band s was critical for subsequent data analysis. To quantify structural parameters, such as sarcomere length, sarcomeric band length, and Z-line thickness, ImageJ 1.53a was employed as a reliable tool.

### Western immunoblotting

WT and *Dsg2*^mut/mut^ LV and RV free wall were lysed in RIPA buffer containing 1:100 protease and phosphatase inhibitors (Sigma; Cat. No. P1860 and P0044, respectively). Protein concentrations were then measured using a BCA protein assay kit (Pierce, Cat. No. 23225). 15 μg of protein from each of the samples were subjected to electrophoresis in 12% SDS-PAGE gel at 100 V. Next, the proteins were transferred to nitrocellulose membranes using the iBlot^TM^ 2 dry blotting system at 20 V for 7 min at room temperature. The membranes were blocked with iBind^TM^ flex FD solution for 30 min at room temperature with gentle agitation. Next, the membranes were placed in the iBind^TM^ flex western device and probed with primary antibodies: rabbit polyclonal anti-GAPDH (Santa Cruz (F L-335): sc-25778) at 1:1000 dilution, rabbit polyclonal anti-MYBPC3-P_ser282_ (custom-made, gift from Dr. Sakthivel Sadayappan, University of Cincinnati) at 1:2000 dilution; mouse monoclonal anti-MYBPC3 (Santa Cruz (E-7): sc-137180) at 1:400 dilution; rabbit polyclonal anti-cTnI-P_ser23/24_ (Cell Signaling; Cat #4004) at 1:200 dilution; mouse monoclonal anti-cTnI total (TI-1, Developmental Studies Hybridoma Bank, University of Iowa) at 1:400 dilution, and rabbit polyclonal anti-α-Actinin 2 (GeneTex #GTX103219) at 1:500 dilution. The secondary antibodies were goat anti-rabbit (IRDye 680RD; LI-CORTM) at 1:1000 dilution and goat anti-mouse (IRDye 800CW; LI-CORTM) at 1:1000 dilution. Both primary and secondary antibodies were prepared in iBind^TM^ flex FD solution. The membranes were washed three times (5 min each) with TBS with 0.05% Tween^TM^ 20. Lastly, the membranes were imaged using the Odyssey^TM^ CLx imaging system (LI-COR^TM^) and quantified using the ImageStudio version 4 (LI-COR^TM^).

### Myofilament and myofibrillar alignment analysis via electron microscopy (EM)

Surrogates for myofilament and myofibrillar disarray were performed as described previously (31). Myofilament disarray was quantified using width and angle sigma, corresponding with lattice and longitudinal myofilament disarray, respectively. Small 70 x 70 pixel kernels were sampled throughout the EM image such that only myofilaments were selected. At least 10 such kernels were selected. For each kernel, two dimensional fast Fourier transforms were calculated, and magnitude images were rearranged such that that zero-frequency term was centered. The resulting spectra were then summed for all kernels. The “equator” in the Fourier domain was manually identified and the image rotated. Numerical radial and angular integrals were quantified and fit with a cubic spline. The resulting pattern took the form of J0 Bessel function, and the second maxima from the center was used for determination of the width. Myofibrillar disarray was quantified also with width and angle sigma, but at lower resolution. The entire image was fast Fourier transformed, magnitude images rotated by manual selection of the equator, and radial and angular integrals quantified with 0.01 degree and 0.01-pixel resolution, respectively. The intensity corresponding with fibril alignment, somewhat analogous to the (1,0) intensity on small angle x-ray diffraction, was isolated and fit to a univariate Gaussian. The variance of each of the Gaussians in the angular and radial directions were taken to be angle and width sigma, respectively. For each image, sarcomere length was also determined, and the distance from the center of the power spectra was calculated, corresponding to inter-myofibrillar spacing.

### Cardiac muscle bundle preparations (CMBs)

Permeabilized LV and RV were prepared as previously described (32). Excised WT and *Dsg2*^mut/mut^ hearts were cut-open along the septum to expose the ventricles to 1% Triton X-100 (v:v) in a pCa 8 relaxing solution (150mM ionic strength, 2.5mM MgATP^2−^, pH 7). CMBs were incubated for 5 h in Triton X-100 permeabilizing solution at 4°C. Next, the permeabilizing solution was removed and a relaxing pCa 8 solution containing 51% glycerol (v:v) was added, and CMBs were stored at −20°C. For muscle mechanics, strips of permeabilized LV and RV free walls were isolated and dissected, attached to aluminum T-clips, and end compliance was minimized by chemically fixing the tissue ends using 1% glutaraldehyde. Sarcomere length (SL) was set to 2.1 μm using HeNe laser diffraction at pCa 8.

### Mechanics of CMBs contraction

Ca^2+^ solutions were calculated utilizing the pCa Calculator (33) containing 20 mM 3-[N-morpholino] propanesulfonic acid (MOPS), 7 mM ethylene glycol-bis(2-aminoethylether)-N,N,N′,N′-tetra acetic acid (EGTA), 15 mM phosphocreatine (CrP), 15 units mL−1 creatine phosphokinase (CPK), 2.5 mM MgATP^2−^, 1 mM free Mg^2+^, ionic strength maintained constant at 150mM by adding KPr, varying [Ca^2+^], pH 7.0. The anion of choice for the pCa solutions was propionate (e.g., CaPr_2_, MgPr_2_ and KPr). All solutions contained 3% Dextran T-500 to bring the myofilament lattice spacing closely to the physiological levels (12). The solutions were made at room temperature (21°C), and experiments were performed at 30°C.

The Ca^2+^-dependence isometric steady-state force levels were measured by a force transducer (Aurora Scientific Inc. Model 403A) was obtained after exposing the cardiac muscle to a series of Ca^2+^ solutions ranging from pCa 8.0 to 4.5 at 30 °C. Muscle length was controlled by a high-speed servomotor (Aurora Scientific Inc. Model 322C). Both pCa_50_ and *n*_Hill_ were estimated by fitting the data using a 2-parameter Hill equation as described (34). After isometric force reached steady state in each pCa solution, mechanics of cardiac muscle contraction were measured as previously described (32, 35). Kinetics of tension redevelopment (*k*_TR_) was obtained by rapidly shortening the cardiac muscle by 19% of its initial length (L_0_) for 20 ms followed by rapid re-stretch (25% L_0_). Then, the cardiac muscle preparation was returned back to L_0_. The apparent rate constant *k* was obtained from each tension recovery time course as previously described (32, 35). Sinusoidal stiffness (SS) measurements were recorded as previously described (32, 35). Briefly, CMBs were oscillated in 0.2% L_0_ (peak-to-peak) at 100 Hz with a sampling rate of 1 kHz. All SS data were analyzed using R Studio and fit with a 4-parameter Hill equation (32, 35).

### Mathematical modeling of kinetics of tension redevelopment

MatLab was used to model the relationship between isometric steady-state force and *k*_TR_ using the 3-state model of muscle contraction and estimate the following parameters: attachment rate (*f*), detachment rate (*g*), and *k*_OFF_ as previously described (12). *k*_ON_ was held constant according to measurements obtained experimentally using WT cardiac muscle samples (36).

### SRX and DRX measurements

Papillary and trabeculae from RV and LV free wall were incubated in a permeabilizing solution (100 mM NaCl, 8 mM MgCl_2_, 5 mM EGTA, 3 mM NaN_3_, 5 mM K_2_HPO_4_, 5 mM KH_2_PO_4_, 5 mM ATP, 20 mM BDM, 1 mM DTT, 0.1 % Triton-X 100, and pH 6.8) at 4°C for 4-6 hours. Then, samples were incubated overnight at 4°C in glycerinating solution and stored in this solution at -20°C. Each sample is then pinned in place on a custom-made glass slide with a coverslip on top to form a chamber. The chamber is filled with 120 ul of skinning solution for 10 mins and washed out with 120 ul Glycerinating solution (120 mM K acetate (Kac), 5 mM Mg acetate_2_ (MgAc_2_), 2.5 mM K_2_HPO_4_, 2.5 mM KH_2_PO_4_, 50 mM MOPS, 5 mM ATP, 20 mM BDM, 2 mM DTT, 50% v/v Glycerol, and pH 6.8) and kept on ice before imaging. Each slide is incubated with 120ul of Rigor buffer (120 mM Kac, 5 mM MgAc_2_, 2.5 mM K_2_HPO_4_, 2.5 mM KH_2_PO_4_, 50 mM MOPS, 2 mM DTT, and pH 6.8) at room temperature for 5 min twice (10 mins in total) before mounting onto the microscope. The pre-set computer program can then be initiated and 120ul of MantATP buffer (250 μM MantATP in Rigor Buffer solution) added to the chamber immediately. This program captures an image every 4 seconds for 10 minutes. After checking the MantATP load phase has successfully increased fluorescence of the sample, the imaging program can be initiated once again and 120ul of ATP chase buffer (120 mM KAc, 5 mM MgAc_2_, 2.5 mM K_2_HPO_4_, 2.5 mM KH_2_PO_4_, 50 mM MOPS, 4 mM ATP, 2 mM DTT, and pH 6.8) pipetted immediately into the chamber. This collects another 10 minutes of images of fluorescence decay. The percent approximation for myosin heads occupying the DRX state was calculated using 0.64 as the correction factor the percent approximation for myosin heads occupying the SRX state was then calculated as 100 – DRX. The videos collected were analyzed using a combination of ImageJ (FIJI), excel, and Prism, and then plotted and fit to a two-phase association curve in Prism.

### Single cardiomyocyte mechanics

Permeabilized cardiomyocytes were isolated as described previously. Briefly, 5-10 mg of flash frozen cardiac tissue was incubated in 0°C isolation buffer (5.55mM Na_2_ATP, 7.11 mM MgCl_2_, 2 mM EGTA, 108.01 mM KCl, 8.91 KOH, 10 mM Imidazol, 10mM DTT) with 0.3% Triton X-100 in the presence of protease (Sigma-Aldrich, MO) and phosphatase inhibitors (PhosSTOP, Roche, Germany) and homogenized at 7500 rpm (OMNI Digital Programmable Homogenizer, Kennesaw, GA). Following homogenization, the sample was incubated in isolation buffer with Triton X-100 for 20 min at 4°C with gentle agitation. The resulting pellet was washed in isolation buffer without Triton X-100. Isolated cardiomyocytes were identified and attached to a force transducer-length controller (Aurora Scientific, Canada) using ultraviolet-activated adhesive (Norland, NJ) in room temperature relaxation buffer 5.95 mM Na2ATP, 6.41 mM MgCl2,10 mM EGTA, 100 mM BES, 10 mM CrP, 50.25 mM KPr, protease inhibitor (Sigma-Aldrich, St. Louis, MO), 1 mM DTT). Sarcomere length (SL) was measured by Fourier transformation (IPX-VGA210, Imperx, FL) and adjusted by micro-manipulators (Siskiyou, CA). All permeabilized single cardiomyocyte experiments were performed at 2.1 µm sarcomere length, in the presence of 3% Dextran T-500 and at 30 °C.

Isometric force Ca^2+^ curves were measured in Ca^2+^ activation buffer ranging from 0.0 to 46.8 µM ((46.8 μM solution composition: 5.95 mM Na_2_ATP, 6.20 mM MgCl_2_, 10 mM Ca^2+^-EGTA, 100 mM BES, 10 mM CrP, 29.98 mM KPr, protease inhibitor (Sigma-Aldrich, St. Louis, MO), 1 mM DTT). Force was calculated by dividing measured force by myocyte cross sectional area, approximated by CSA=π/4 a^2^, where a is the measured myocyte diameter. The rate of isometric tension redevelopment k_TR_ was measured at maximum activation (46.8 µM) by fitting the exponential force recovery following a 20% reduction in SL for 20 ms and re-stretch to a mono-exponential function. In addition, at maximum activation a 2% cell length step response (<0.15% sarcomere length change) was acquired and measured for 1 second at 2000 Hz. The resulting force transient followed a classic three phase process, denoted as the A process, B process, and C process, A corresponding to stiffness of the myocyte and B and C to myosin cross-bridge attachment and detachment, respectively, described elsewhere in detail (37). This force transient was normalized such that maximum activated force was set to 0, and the peak following the A process normalized to 1. The force transient was fit to the equation *F̅*(*t*) = *P*_1_ + *P*_2_*e*^−2π*b t*^ − *P*_3_*e*^−2π*ct*^, from which myosin attachment (2π*b*) and detachment (2π*c*) rates were acquired.

### Small-angle X-ray diffraction

Equatorial X-ray diffraction patterns were collected from strips of permeabilized LV and free walls using the small-angle instrument on the Biological Small Angle X-ray Solution Scattering and High-Pressure Biology Beamline (7A) at Cornell High Energy Synchrotron Source (CHESS). The X-ray beam was collimated to ∼0.25 × 0.25 mm. The sample-to-detector distance was ∼1.7 m and the X-ray wavelength was 0.103 nm. CMBs were dissected (∼200 μm diameter, ∼1.5 - 2 mm long) and permeabilized with 1% Triton X-100 (v:v) in a pCa 8 relaxing solution (described above) overnight. Sarcomere length was adjusted by laser diffraction using a 4-mW HeNe laser. Diffraction patterns were collected at sarcomere lengths of 2.1 μm and in pCa 8 relaxing solution at 30 °C. X-ray exposures were 1 s at an incident flux of ∼10^12^ photons per second, and the patterns were collected on the Eiger 4M detector (Dectris, Switzerland. The intensity ratio and lattice spacing were extracted from X-ray diffraction patterns using the “Equator” routine within the MuscleX software package, developed at BioCAT (38).

### Statistical Analyses

All experiments utilized age-matched males and female mice. Data are presented as mean ± SE. Student’s t-test or two-way ANOVA followed by appropriate post-hoc analysis to determine differences between measured variables, and statistical significance set at P < 0.05. All n-values and the statistical test performed are indicated within each figure legend.

---

**EXTENDED DATA FIGURE 1.**
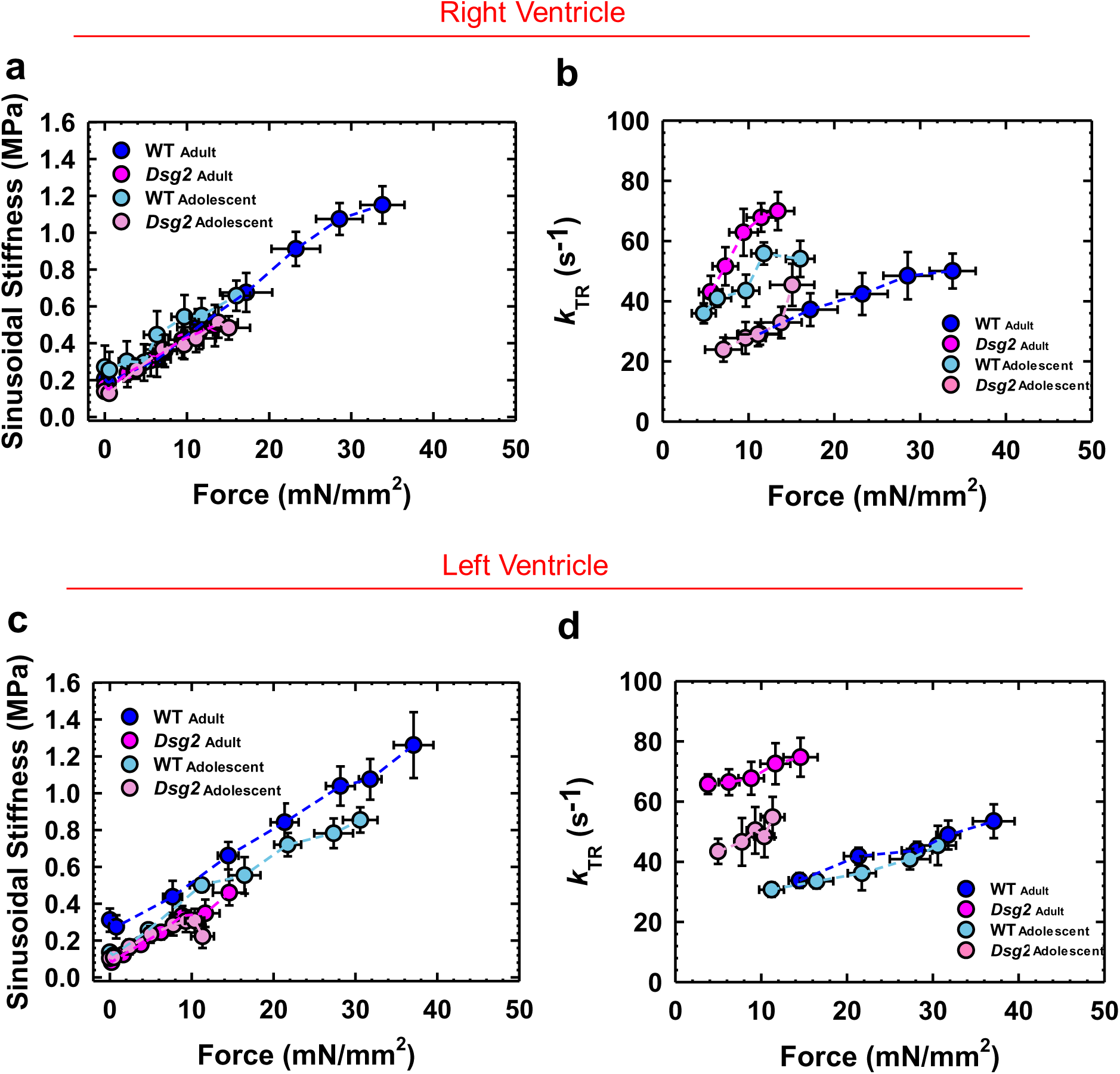
Extended figure legend 1. Effects of the DSG2 deficiency on sinusoidal stiffness (a and c) and kinetics of tension redevelopment (*k*_TR_) (b and d). Steady-state isometric force levels normalized to the cross-sectional area of the cardiac muscle preparations. Data are shown as mean ± S.E. Number of independent animals per group: adolescent WT: 5-7 and adult WT:6-8; adolescent *Dsg2*^mut/mut^: 4-7 and adult *Dsg2*^mut/mut^: 6-9.

**EXTENDED DATA FIGURE 2.**
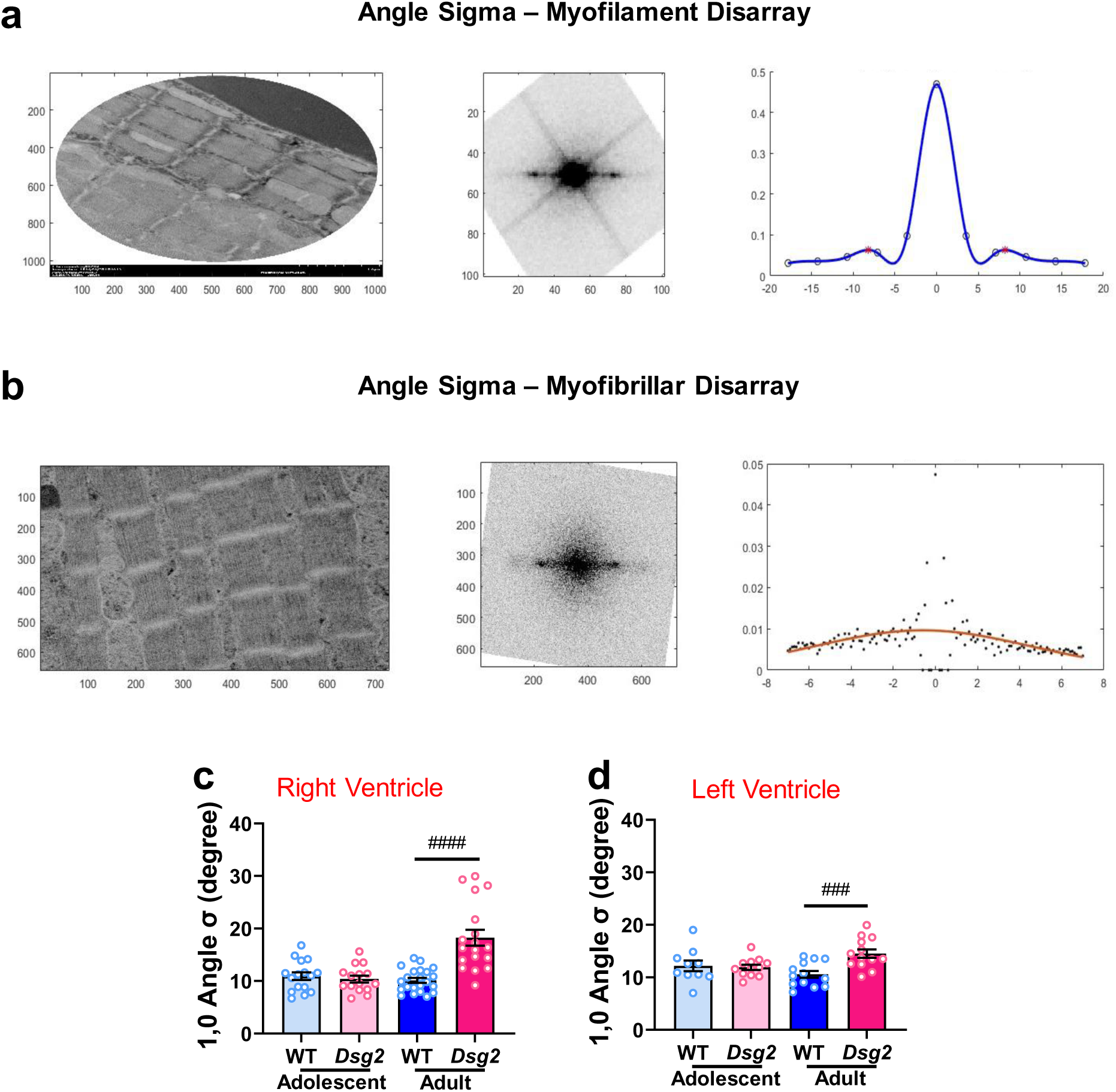
Extended figure legend 2. Myofilament and myofibrillar disarray analysis. **a)** Representative EM image analysis workflow for myofilament disarray: representative EM image ◊ Power Spectrum ◊ Spline Interpolation/First Order Bessel *Function* (blue) to quantify angle σ of the angular integral of the (1,0) intensity from the power spectrum. **b)** Representative EM image analysis workflow for myofibrillar disarray: representative EM image ◊ Power Spectrum ◊ Gaussian fit (orange) to quantify angle σ of the angular integral of the (1,0) intensity from the power spectrum. The angular standard deviation of 1,0 equatorial reflections (angle σ) from longitudinal EM images of RV **(c)** and LV **(d)** of adolescent and adult WT and *Dsg2*^mut/mut^. Data are shown as mean ± S.E. Statistical significance was assessed by two-way ANOVA followed by Bonferroni’s multiple comparisons test. ^###^ P < 0.001 for *Dsg2*^mut/mut^ vs. WT mice. n = 3 independent animals per group were utilized. Number of EM images: Adolescent RV, WT n = 16 and *Dsg2*^mut/mut^ n = 15; Adolescent LV, WT n = 10 and *Dsg2*^mut/mut^ n = 12; Adult RV, WT n = 22 and *Dsg2*^mut/mut^ n = 18; adult LV, WT n = 13 and *Dsg2*^mut/mut^ n = 13.

**EXTENDED FIGURE 3.**
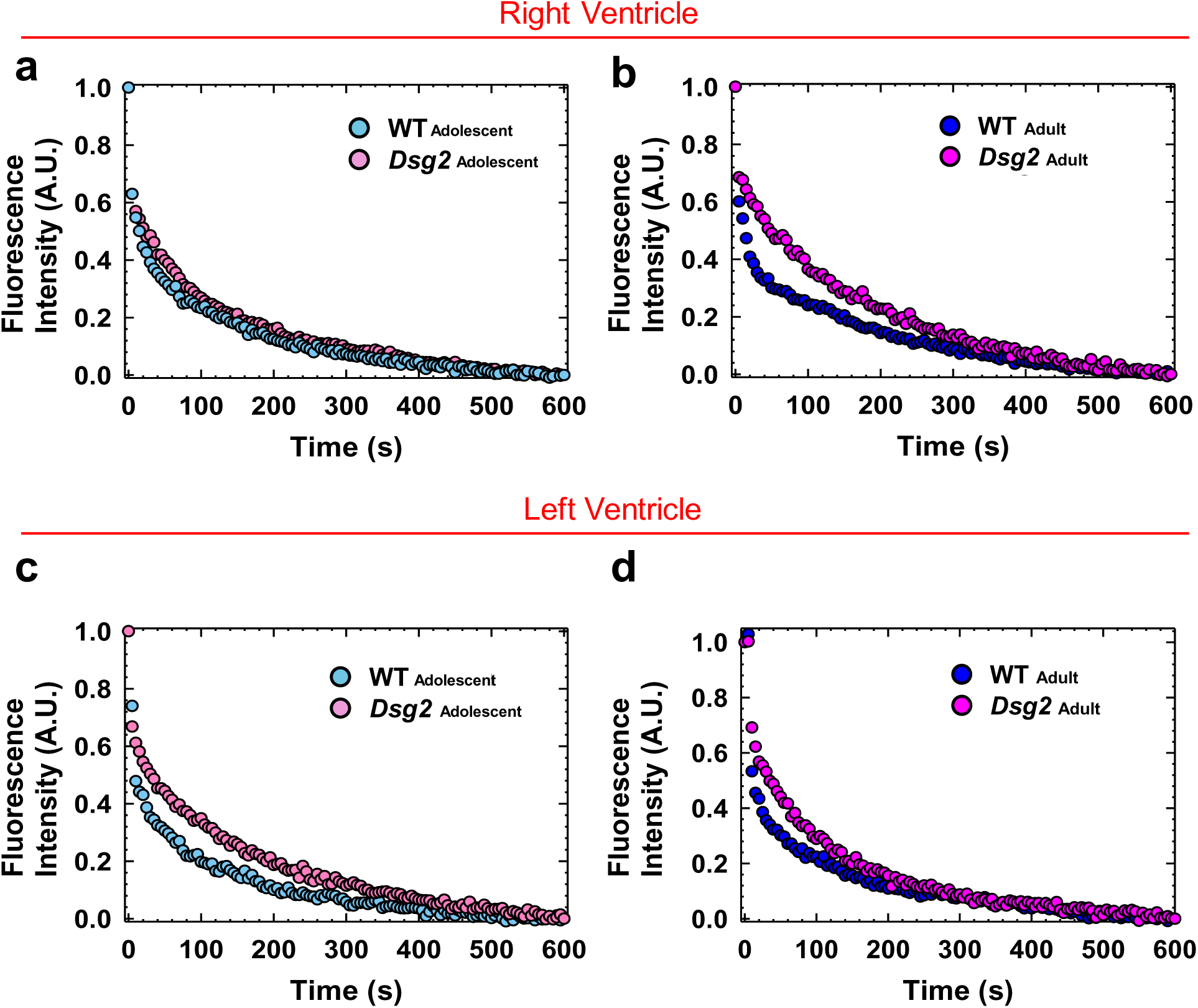
Extended figure legend 3. Effects of the DSG2 deficiency on the fraction of DRX and SRX myosin populations. Representative Mant-ATP fluorescence decay assay performed in permeabilized RV **(a-b)** and LV **(c-d)** free wall of adolescent and *adult* WT and *Dsg2*^mut/mut^ mice.

**SUPPLEMENTARY FIGURE 1.**
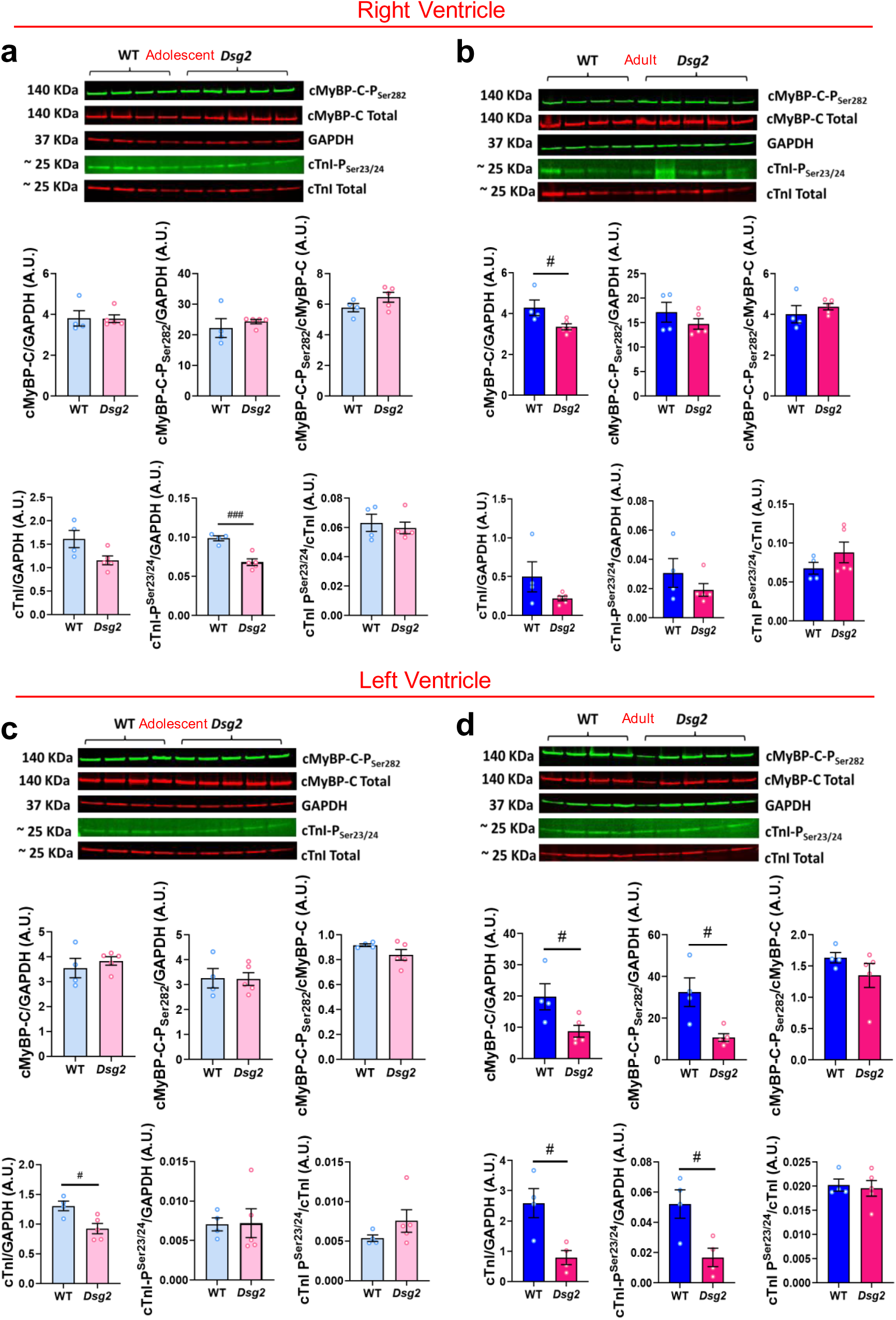
Supplementary figure legend 1. Impact of DSG2 deficiency on myofilament protein levels and phosphorylation status. Cropped images of western immunoblotting membranes incubated with anti-cMyBP-C, -cMyBP-C-P_ser282_, -GAPDH, -cTnI, and -cTnI-P_ser23/24_ antibodies, and their respective immunoblot quantification. Tissue lysates were prepared from RV **(a-b)** and LV **(c-d)** free walls of adolescent and adult WT and *Dsg2*^mut/mut^ mice. Data are shown as mean ± SE. Statistical significance was assessed by unpaired Student t-test between adolescent *Dsg2*^mut/mut^ vs. WT mice; ^#^ P < 0.05 and ^##^ P < 0.01. Number of biological replicates: WT n = 4 and *Dsg2*^mut/mut^ n = 5.

**SUPPLEMENTARY TABLE 1.** Echocardiographic and morphometric indices from adolescent WT and *Dsg2*^mut/mut^ mice. IVRT, interventricular relaxation time; IVCT, interventricular contraction time; MV A/E, mitral valve early peak flow velocity/atrial peak flow velocity; IVSd, interventricular septal end-diastolic volume; IVSs, interventricular septal end-systolic volume; LVPWd, left ventricular posterior wall end diastole; LVPWs, left ventricular posterior wall end systole; LVAWd, left ventricular anterior wall end diastole; LVAWs, left ventricular anterior wall end systole; SV, stroke volume; CO, cardiac output; FS, fractional shortening; EF, ejection fraction; RWT, relative wall thickness; HW, heart weight; and BW, body weight. Data presented as mean ± SE, ^#^ P < 0.05 and ^###^ P < 0.001 for adolescent *Dsg2*^mut/mut^ vs. WT mice using unpaired Student’s *t*-test. ECHO, WT n = 11 and *Dsg2*^mut/mut^ n = 7-9; and morphometric, WT n = 11-15 and *Dsg2*^mut/mut^ n = 7-20.

| Parameters | WT | *Dsg2*<sup>mut/mut</sup> |
| --- | --- | --- |
| **Echocardiographic** |  |  |
|  | Adolescent |  |
| IVRT (ms) | 14.39 ± 0.64 | 15.41 ± 0.86 |
| IVCT (ms) | 12.95 ± 1.05 | 14.72 ± 1.62 |
| MV A/E (A.U.) | 1.97 ± 0.07 | 1.61 ± 0.06<sup>###</sup> |
| IVSd (mm) | 0.65 ± 0.03 | 0.57 ± 0.03 |
| IVSs (mm) | 0.72 ± 0.05 | 0.65 ± 0.04 |
| LVPWd (mm) | 0.58 ± 0.05 | 0.54 ± 0.01 |
| LVPWs (mm) | 0.80 ± 0.05 | 0.71 ± 0.02 |
| LVAWd (mm) | 0.78 ± 0.04 | 0.68 ± 0.03 |
| LVAWs (mm) | 1.10 ± 0.03 | 0.88 ± 0.05<sup>###</sup> |
| SV (uL) | 31.64 ± 1.91 | 26.85 ± 1.57 |
| CO (mL/min) | 13.56 ± 0.77 | 10.99 ± 1.03 |
| FS (%) | 27.57 ± 1.61 | 22.07 ± 1.79<sup>#</sup> |
| EF (%) | 54.02 ± 2.63 | 44.61 ± 3.23<sup>#</sup> |
| Dyskinesia (%) | 9 (n = 1 / 11) | 42 (n = 6 / 14) |
| Hypokinesia (%) | 9 (n = 1 / 11) | 57 (n = 8 / 14) |
| Hyperkinesia (%) | 0 (n = 0 / 11) | 14 (n = 2 / 14) |
| **Morphometric** |  |  |
|  | Adolescent |  |
| RWT (mm) | 0.31 ± 0.01 | 0.29 ± 0.01 |
| HW (mg) | 74.89 ± 4.31 | 72.45 ± 1.39 |
| BW (g) | 15.70 ± 0.91 | 13.64 ± 0.34 |
| HW / BW (mg/g) | 4.78 ± 0.09 | 5.36 ± 0.15<sup>#</sup> |

**SUPPLEMENTARY TABLE 2.** Contractile parameters measured in adolescent and adult WT and *Dsg2*^mut/mut^ mouse permeabilized left and right ventricular free wall as depicted in. Figures 2. F_max_, maximum steady-state isometric force; pCa_50_, Ca^2+^ concentration needed to reach 50% of the steady-state isometric maximum force; *n*_Hill_, cooperativity of thin filament activation; *k*_TR_ max, maximum rate of tension redevelopment; SS_pass_, passive steady-state sinusoidal stiffness values obtained at pCa 8 (resting condition); SS_max_, maximum steady-state sinusoidal stiffness; CMBs, cardiac muscle preparations. Data are shown as mean ± S.E. Statistical significance was assessed by two-way ANOVA followed by Tukey’s or Bonferroni’s multiple comparison test. * P < 0.05, ** P < 0.01, and *** P < 0.001 within the same genotype; ^#^ P < 0.05 and ^###^ P < 0.001 between genotypes.

|  | Right Ventricle |  |  |  |
| --- | --- | --- | --- | --- |
|  | WT |  | *Dsg2*<sup>mut/mut</sup> |  |
|  | Adolescent | Adult | Adolescent | Adult |
| **F<sub>max</sub>** (mN/mm<sup>2</sup>) | 16.01 ± 1.66 | 33.78 ± 2.67*** | 15.08 ± 2.58 | 13.40 ± 1.91### |
| **FpCa<sub>50</sub>** | 5.61 ± 0.09 | 5.67 ± 0.06 | 5.77 ± 0.06 | 5.69 ± 0.05 |
| **Fn<sub>Hill</sub>** | 1.80 ± 0.20 | 1.87 ± 0.10 | 1.93 ± 0.15 | 1.82 ± 0.15 |
| **k<sub>TR max</sub>** (s<sup>-1</sup>) | 54.09 ± 6.03 | 50.06 ± 5.77 | 45.44 ± 7.02 | 69.99 ± 6.34* |
| **SS<sub>pass</sub>** (MPa) | 0.27 ± 0.11 | 0.20 ± 0.06 | 0.13 ± 0.03 | 0.16 ± 0.05 |
| **SS<sub>max</sub>** (MPa) | 0.65 ± 0.08 | 1.15 ± 0.10** | 0.48 ± 0.06 | 0.49 ± 0.11### |
| **#CMBs** | 5-7 | 7-8 | 7 | 6-9 |

**Supplementary table 2. Contractile parameters measured in adolescent and adult WT and *Dsg2<sup>mut/mut</sup>* mouse permeabilized left and right ventricular free wall as depicted in Figures 2.**
|  | Left Ventricle |  |  |  |
| --- | --- | --- | --- | --- |
|  | WT |  | *Dsg2*<sup>mut/mut</sup> |  |
|  | Adolescent | Adult | Adolescent | Adult |
| **F<sub>max</sub>** (mN/mm<sup>2</sup>) | 30.58 ± 2.11 | 37.10 ± 2.42** | 11.32 ± 1.39### | 14.57 ± 1.99### |
| **FpCa<sub>50</sub>** | 5.69 ± 0.03 | 5.75 ± 0.04 | 5.81 ± 0.08<sup>#</sup> | 5.57 ± 0.03***<sup>#</sup> |
| **Fn<sub>Hill</sub>** | 1.91 ± 0.11 | 1.83 ± 0.12 | 2.64 ± 0.42<sup>#</sup> | 1.83 ± 0.09* |
| **k<sub>TR max</sub>** (s<sup>-1</sup>) | 45.44 ± 6.54 | 53.50 ± 5.60 | 54.82 ± 6.76 | 74.77 ± 6.47 |
| **SS<sub>pass</sub>** (MPa) | 0.13 ± 0.02 | 0.31 ± 0.06 | 0.11 ± 0.03 | 0.10 ± 0.01 |
| **SS<sub>max</sub>** (MPa) | 0.85 ± 0.06 | 1.26 ± 0.17** | 0.24 ± 0.07### | 0.45 ± 0.06### |
| **#CMBs** | 5-7 | 7-8 | 4-7 | 7-8 |

**SUPPLEMENTARY TABLE 3.**
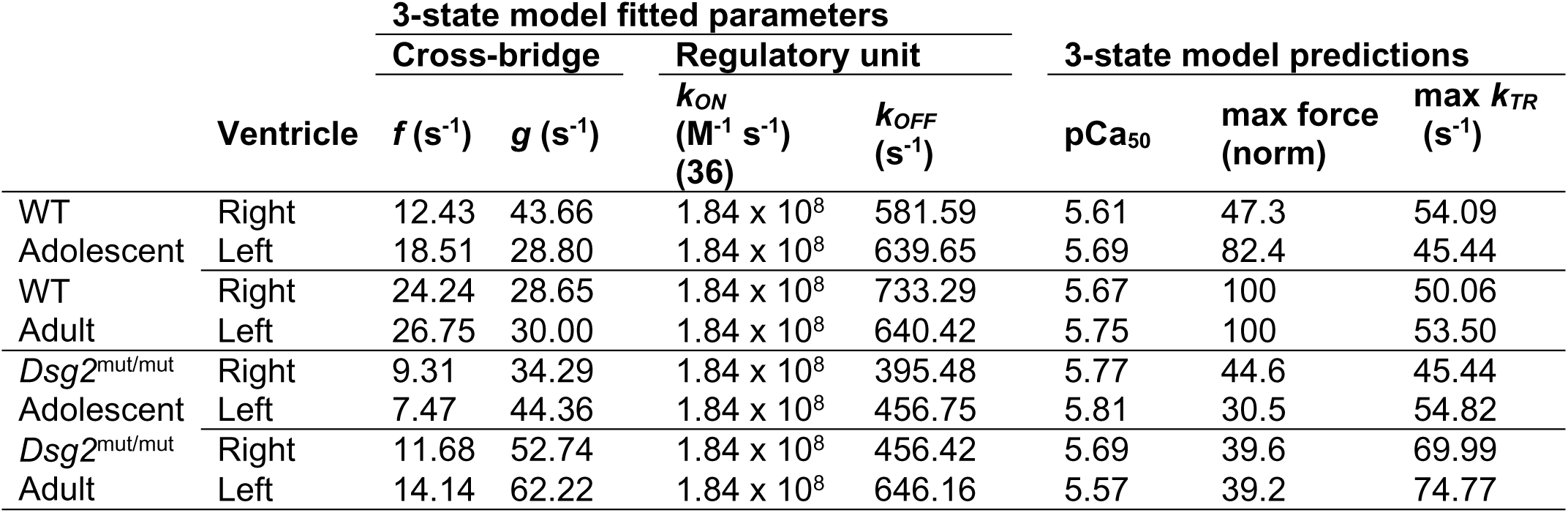
Optimized parameter predictions and estimates from the three-state model for steady-state isometric force-*k*_TR_ data depicted in. Fig. 2. All best-fit parameter estimates for *f* (attachment rate), *g* (detachment rate), and *k*_OFF_ (Ca^2+^ dissociation rate) were obtained in MatLab by using the simplex method for the steady-state isometric force-*k*_TR_ data shown in Fig. 2. The simplex method considers the values of myofilament Ca^2+^-sensitivity (pCa_50_) from steady-state isometric force measurements as reported in Fig. 2; values for *k*_ON_ were derived from measurements reported by Pinto et al. (36), and maximum steady-state isometric force at WT adult assumed to be 1.0 for each dataset. These parameter sets were used to fit and illustrate relations between *k*_TR_ and steady-state isometric force in Fig. 2 (dashed lines in Fig. 2e and j).

## Acknowledgements

This work was supported by an American Heart Association Career Development Award [19CDA34760185 (to SPC) and a Florida State University Research Foundation Award (to SPC). X-Ray diffraction experiments were supported by grant P30 GM138395 from the National Institute of General Medical Sciences of the National Institutes of Health and were conducted at the Center for High-Energy X-ray Sciences (CHEXS), supported by the National Science Foundation (BIO, ENG and MPS Directorates) under award DMR-1829070, and at the Macromolecular Diffraction at CHESS (MacCHESS) facility, supported by award 1-P30--GM124166--01A1 from the National Institute of General Medical Sciences, National Institutes of Health, and by New York State’s Empire State Development Corporation (NYSTAR). The British Heart Foundation Doctoral training centre in Oxford supported FP. CT is a Sir Henry Dale Fellowship with code (Grant 222567/Z/21/Z).

